# Motor cortical influence relies on task-specific activity covariation

**DOI:** 10.1101/2022.02.09.479479

**Authors:** Claire L. Warriner, Samaher Fageiry, Shreya Saxena, Rui M. Costa, Andrew Miri

**Author notes:** Correspondence (A.M.).

## Abstract

During limb movement, spinal circuits facilitate the alternating activation of antagonistic flexor and extensor muscles. Yet antagonist cocontraction is often required to stabilize joints, like when loads are handled. Previous results suggest that these different muscle activation patterns are mediated by separate flexion- and extension-related motor cortical output populations, while others suggest recruitment of task-specific populations. To distinguish between hypotheses, we developed a paradigm in which mice toggle between forelimb tasks requiring antagonist alternation or cocontraction and measured activity in motor cortical layer 5b. Our results conformed to neither hypothesis: consistent flexion- and extension-related activity was not observed across tasks, and no task-specific populations were observed. Instead, activity covariation among motor cortical neurons dramatically changed between tasks, thereby altering the relation between neural and muscle activity. This was also observed specifically for corticospinal neurons. Collectively, our findings indicate that motor cortex drives different muscle activation patterns via task-specific activity covariation.

**HIGHLIGHTS:**

- Mice perform two forelimb tasks involving distinct antagonist muscle activity in a novel paradigm
- L5b motor cortical neurons are not organized by task-specific activity
- L5b motor cortical neurons do not encode muscle activity consistently across tasks
- Task-specific muscle activity is driven by a change in motor cortical activity covariation

**eTOC BLURB:** Warriner et al. simultaneously measured muscle and motor cortical activity in mouse during antagonist forelimb muscle alternation and cocontraction, revealing that these distinct muscle activation patterns are not driven through consistent flexion and extension programs nor through the activity of discrete, task-specific neuronal subsets. Instead, distinct patterns involve task-specific changes in firing pattern covariation among layer 5b neurons, and corticospinal neurons in particular, which change their relationship to muscle activity across tasks.

## INTRODUCTION

Whether running from a hungry cheetah or maintaining a strong grip on a writhing antelope, mammals depend on the ability to control their limb movement and stiffness for survival. Antagonistic limb muscles exert opposing forces on joints, and distinct temporal patterns of their activation drive different motor outputs according to context. Joint rotations involve the inversely related activation (alternation) of antagonist muscles. However, motor behaviors often require joints be stabilized rather than rotated, necessitating the simultaneous contraction of antagonist muscles. This cocontraction is observed, for example, when limb position is maintained against a load (Milner, 2002; Smith, 1981), during the anticipation of a load as when catching a ball (Lacquaniti and Maioli, 1987), and during the early stages of motor skill learning (Gribble et al., 2003; Huang et al., 2012; Thoroughman and Shadmehr, 1999).

Antagonist alternation and cocontraction involve different patterns of spinal circuit engagement, and motor cortex has been implicated in each. To promote alternation during joint rotation, sensory feedback from muscles activates spinal circuits that inhibit the motor neurons of antagonist muscles (Eccles and Lundberg, 1958; Jankowska, 1992). During cocontraction, this reciprocal inhibition is reduced via inhibition of Ia reciprocal inhibitory interneurons and an increase in presynaptic inhibition at sensory afferent terminals (Nielsen and Kagamihara, 1992, 1993; Perez et al., 2007). During voluntary cocontraction, these changes appear to be under supraspinal control (Lévénez et al., 2008; Nielsen and Kagamihara, 1992). Corticospinal neurons (CSNs) contact spinal interneurons that influence antagonist motor neuron activity, both directly, via Ia interneurons and Renshaw cells (Jankowska et al., 1976; Mazzocchio et al., 1994) and indirectly, via presynaptic inhibitory (GABApre) interneurons (Russ et al., 2013). In addition, the strength of reciprocal inhibition can be modified through training in humans and rats (Chen et al., 2006a; Perez et al., 2007) and spinal reflex gain can be regulated by corticospinal neurons (Chen et al., 2006b). However, it remains unclear how motor cortical output promotes these different forms of spinal activity and the modulation of reciprocal inhibition during voluntary movement.

Two potential frameworks for the motor cortical mediation of these different spinal activity modes have been previously proposed. In one view, motor cortex has neuronal populations whose activity is specific to either mode (Figure 1A). Among primate layer 5 (L5) primary motor cortical neurons, a subset shows activity modulation in relation to wrist muscle cocontraction but not flexion or extension (Fetz and Cheney, 1987; Humphrey and Reed, 1983). Zones of primary motor cortex that when stimulated induce antagonist cocontraction have also been observed (Humphrey and Reed, 1983). Human functional imaging has shown that the region of cortex activated during cocontraction is only partially overlapping with areas activated during flexion or extension (Johannsen et al., 2001). Support for task-specific populations of corticospinal neurons also comes from human studies that found differences in the effect of transcranial magnetic stimulation on muscle excitability between cocontraction and contraction of each muscle alone (Aimonetti and Nielsen, 2002; Nielsen et al., 1993). The existence of task-specific output populations would align with evidence supporting a task-based cellular organization of motor cortex from studies in monkeys (Graziano et al., 2002, 2005), rats (Brown and Teskey, 2014), and mice (Harrison et al., 2012).

**Figure 1.**
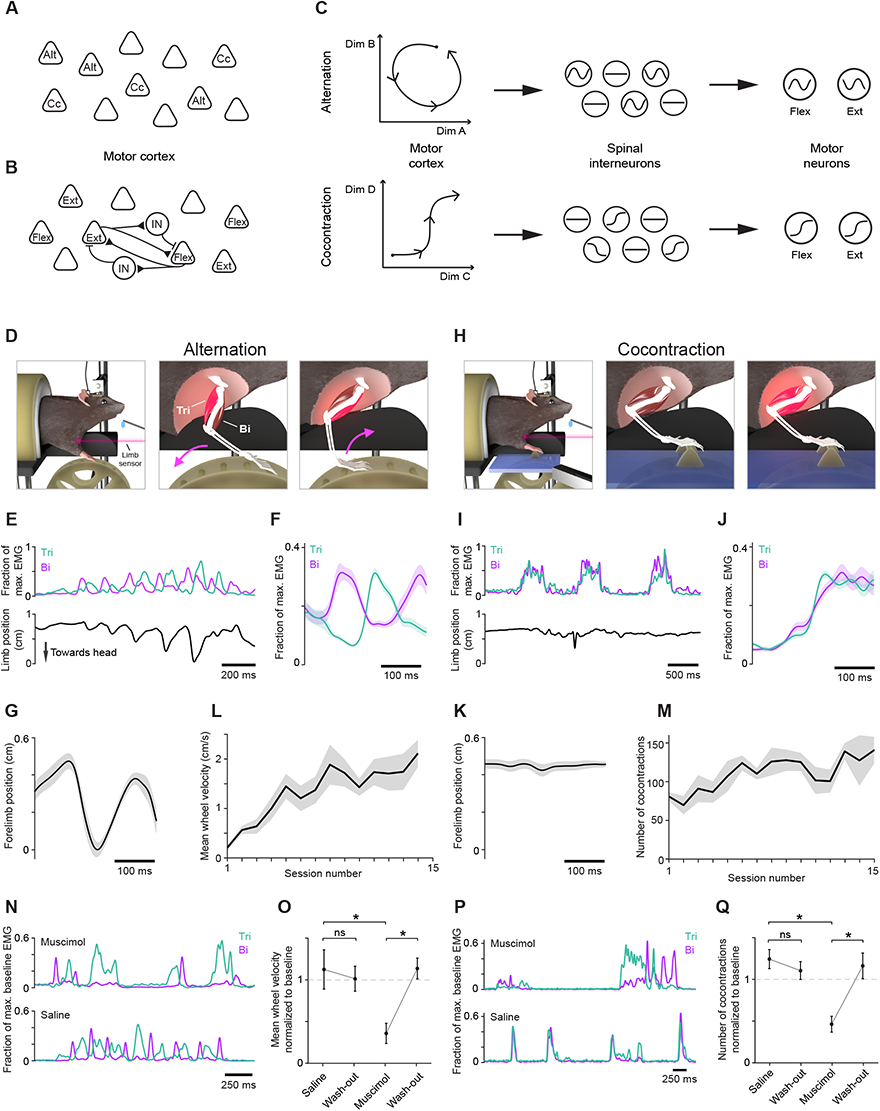
Alternation and cocontraction behavioral paradigm. (A) Motor cortex may use task-specific pyramidal neurons to mediate cocontraction (Cc) and alternation (Alt). (B) Alternately, cocontraction may be produced through the concurrent activation of flexion (Flex) and extension (Ext) related pyramidal neurons, whose activity is gated by interposed interneurons (IN) during tasks not requiring cocontraction. (C) In a third possibility, motor cortical neurons may be active during both tasks and their population-level changes in covariance structure differentially engage spinal interneurons to produce flexor-extensor alternation (top) or cocontraction (below). (D, H) Schematic depicting the alternation (D) and cocontraction (H) tasks and associated muscle activity (bright red) (E-F, I-J) EMG from triceps (Tri) and biceps (Bi) activity during the alternation (E-F) and cocontraction (I-J) tasks. (E) and (I) show example EMG time series with limb position below. (F) and (J) show trial-averaged EMG mean ± SEM (n = 30 trials) from an individual session. (G, K) Trial-averaged limb position ± SEM (n = 30 trials) aligned to triceps onset from alternation (G) and cocontraction (K) epochs from the same time points as shown in (F) and (J), respectively. (L-M) Mean ± SEM (n = 5 mice) of mean wheel velocity during alternation training (L) and number of cocontraction events during cocontraction training (M). Analysis limited to the first 20 minutes of each session due to variation in session length. (N-Q) Muscimol or saline was injected into contralateral CFA and performance was tested 90 minutes later. Wash-out sessions occurred at least 12 hours after injection. Examples of triceps (Tri) and biceps (Bi) activity after injection of muscimol (top) or saline (bottom) during alternation (N) and cocontraction (P). Mean ± SEM (n = 5 mice) in the first 20 minutes of sessions for mean wheel velocity during alternation, normalized to baseline, which is defined as mean performance over the three sessions preceding injection (O; one-tailed paired t-test: saline and wash-out p = 0.642, muscimol and wash-out p = 0.002, saline and muscimol p = 0.023) and cocontraction (Q; saline and wash-out p = 0.766, muscimol and wash-out p = 0.004, saline and muscimol p = 0.004).

In a second view, motor cortex mediates different spinal activity modes through activation of flexion- and extension-related output neurons (Figure 1B). In cat, motor cortical regions in which stimulation drives either flexion or extension of the wrist are synaptically coupled, both directly and via intervening inhibitory interneurons (Capaday et al., 1998; Schneider et al., 2002). When inhibition in one of these regions is blocked, stimulation of the other, which previously evoked contraction of one muscle, instead produces cocontraction (Ethier et al., 2007). These and other findings suggest that cocontraction involves a reduction in intracortical inhibition in motor cortex that promotes simultaneous activation of neurons in flexion-and extension-related populations, leaving downstream regions to mediate the reduction of spinal reciprocal inhibition (Capaday, 2004; Capaday et al., 2013; Matsumura et al., 1991). Such a mechanism relies on a muscle-based cellular organization of motor cortex where cells control activation of specific sets of muscles, in agreement with interpretations of the somatotopy revealed by electrical stimulation (Fritsch and Hitzig, 1870; Leyton and Sherrington, 1917; Penfield and Rasmussen, 1950; Tennant et al., 2011).

Here we also consider a third view based on recent results suggesting that changes in motor cortical influence can be mediated by a change in the covariation of its output activity patterns (Figure 1C; Druckmann and Chklovskii, 2012; Elsayed et al., 2016; Kaufman et al., 2014; Miri et al., 2017). In a neural activity space whose cardinal dimensions each represent the firing rate of an individual neuron, changes in activity covariation are reflected in state variation along different directions. In this third view, neurons in downstream circuits would respond specifically to the strong variation along certain directions specific to cocontraction and, in turn, drive changes in spinal circuit behavior that facilitate cocontraction. Conversely, motor cortical activity along other activity space directions would induce spinal circuit behavior that instead promotes alternation. This view aligns with the emerging idea that motor cortical function is not organized solely by cell groups, but by task-specific population dynamics that can flexibly control a variety of motor behaviors (Perich et al., 2018; Russo et al., 2018, Gallego et al., 2017; Sussillo et al., 2015).

To address how motor cortex mediates tasks comprising different muscle activation patterns, we performed population-level analyses of neural activity in layer 5b (L5b) of motor cortex as mice behaved in a novel behavioral paradigm requiring both antagonist alternation and cocontraction. Counter to the view reflecting a task-based cellular organization, neurons showing task-specific activity were rare and existed on a continuum of task-dependent activity modulation across neurons. The view grounded in a muscle-based cellular organization predicts a consistent relationship between neural and muscle activity during alternation and cocontraction. Instead, we found that neuronal populations do not linearly encode muscle activity consistently across tasks. Moreover, we did not find support for an attenuation of intracortical inhibition that enables coactivation of distinct motor cortical output populations. Consistent with the view based on changes in output activity covariation, we found that activity subspaces were strikingly different between the alternation and cocontraction tasks, both for all L5b neurons, and for corticospinal neurons specifically. Our results indicate that motor cortex mediates tasks comprising different muscle activation patterns through changes in the covariation of its output activity patterns.

## RESULTS

### A behavioral paradigm where mice activate antagonist muscles in distinct patterns

We sought to design a behavioral paradigm that would elicit motor cortex-dependent flexor-extensor alternation and cocontraction in mice in a manner conducive to physiological interrogation. We developed a self-paced paradigm in which head-fixed mice perform two different tasks in distinct epochs in order to receive water rewards while forelimb muscle activity is recorded. In an initial epoch, mice iteratively turn a small wheel with their right forelimb until a cumulative turn distance threshold is reached (Figure 1D, Movie S1), which involves alternating activation of triceps (lateral head; elbow extensor) and biceps (elbow flexor; Figure 1E-G). After achieving a set number of rewards, a small platform supporting a single static rung extends, covering the wheel (Figure 1H, Movie S2). Mice then receive rewards for simultaneously activating right triceps and biceps beyond a threshold duration after a period of muscle quiescence (Figure 1I-J). This generally involved little to no movement, which was verified using a laser displacement sensor trained on the front of the mouse’s forelimb (Figure 1H,I,K). To control for within-session variation in neural activity, behavioral sessions end with an additional epoch of the alternation task (Figure S1A). In this paradigm, mice maintained a hunched, squatting posture, retaining free movement of their forelimbs; they periodically groomed their faces with both forelimbs without noticeable change in posture. The activity of pairs of antagonist muscles at the shoulder, elbow, and wrist was recorded with chronically implanted electromyographic (EMG) electrodes (Figure S1B-E; Akay et al., 2006; Miri et al., 2017).

Training for this paradigm involved behavioral shaping over twice-daily sessions lasting up to 45 minutes. For the first 7 days of training, mice performed only the alternation task. Training for the cocontraction task took place over the subsequent 7 days, following which mice were introduced to the combined task paradigm. During alternation task training, the cumulative turn distance threshold was updated within sessions to stabilize the reward rate and encourage longer bouts of wheel turning. Similarly, the cocontraction activity magnitude and duration thresholds for reward were updated within sessions to motivate stronger and longer cocontraction bouts. To further encourage performance improvement, reward volume depended on performance relative to recent attempts. During initial alternation training, mean wheel velocity increased steadily across sessions (Figure 1L, n = 5). During cocontraction training, rapid task acquisition was generally seen within the first session, followed by more modest gains in subsequent sessions; this was reflected in an increased cocontraction frequency assuming fixed criteria for defining cocontraction events, despite the fluctuating reward thresholds (Knarr et al., 2012; see Methods: EMG Processing and Analysis; Figure 1M, n = 5).

Both the alternation and cocontraction tasks depended on motor cortical activity. After one week of training on either task, the GABA_A_ agonist muscimol (74 nl of 1 ng/nl) or saline was injected into the caudal forelimb area (CFA) of left motor cortex (Tennant et al., 2011), and behavioral performance was tested 90 minutes later. During alternation, muscimol injection reduced mean wheel velocity to 35 ± 12% of baseline (mean ± standard error (SEM), n = 5 mice; Figure 1N-O). This was due to a reduction in both the time spent turning the wheel (53 ± 14%, Figure S1F), which suggested a possible deficit in movement initiation, and in mean wheel velocity measured only while the wheel was turning (60 ± 9%, Figure S1G). Mean triceps and biceps activation during wheel turning were unchanged (Tri: 98 ± 4%, Bi: 95 ± 6%; Figure S1H), indicating that inhibition of motor cortex did not reduce the strength of muscle activation during periods when muscles were active. This lack of reduction in muscle activity paired with the reduction in mean wheel speed during wheel rotation suggest a reduction in the efficacy of motor output, perhaps from reduced muscle coordination. These findings also indicate that performance impairment was not due entirely to muscle weakness or paralysis. Similar effects on mean wheel velocity were seen with bilateral muscimol injection (Figure S1I).

During the cocontraction task, unilateral muscimol injection reduced the number of successful cocontraction events, again defined using fixed criteria for event detection across sessions as in Figure 1M, to 46 ± 9% of baseline (mean ± SEM, n = 5 mice; Figure 1P-Q). This was not due to general muscle weakness but appeared to be at least partly due to a reduced ability to initiate cocontraction events. The fraction of time muscles were activated decreased with muscimol injection (54 ± 7%; Figure S1J), but there was little difference in mean triceps and biceps activation levels during task performance (Tri: 99 ± 7, Bi: 89 ± 7; Figure S1L). We used an index of muscle cocontraction to probe for changes in coordination but found only a subsignificant reduction during periods of muscle activity (Knarr et al., 2012; see Methods: EMG Processing and Analysis; 80 ± 9%; Figure S1K). A similar reduction in cocontraction performance was seen after bilateral injection (Figure S1M).

### Motor cortical activity during antagonist muscle alternation and cocontraction

We recorded neural activity in left CFA during behavioral sessions after mice had been fully trained. Recordings were made using acute insertion of a 32-site silicon probe with electrodes targeted to layer 5b (650-850 µm below pia) where subcerebral projection neurons reside. In each mouse, single unit activity was extracted (Rossant et al., 2016) from recordings made at different sites over 6-8 behavioral sessions (Figure 2A). Recording at multiple sites ensured a broad sampling of CFA, where regions associated with individual muscles or joints overlap extensively (Ayling et al., 2009; Ethier et al., 2007; Rathelot and Strick, 2009).

**Figure 2.**
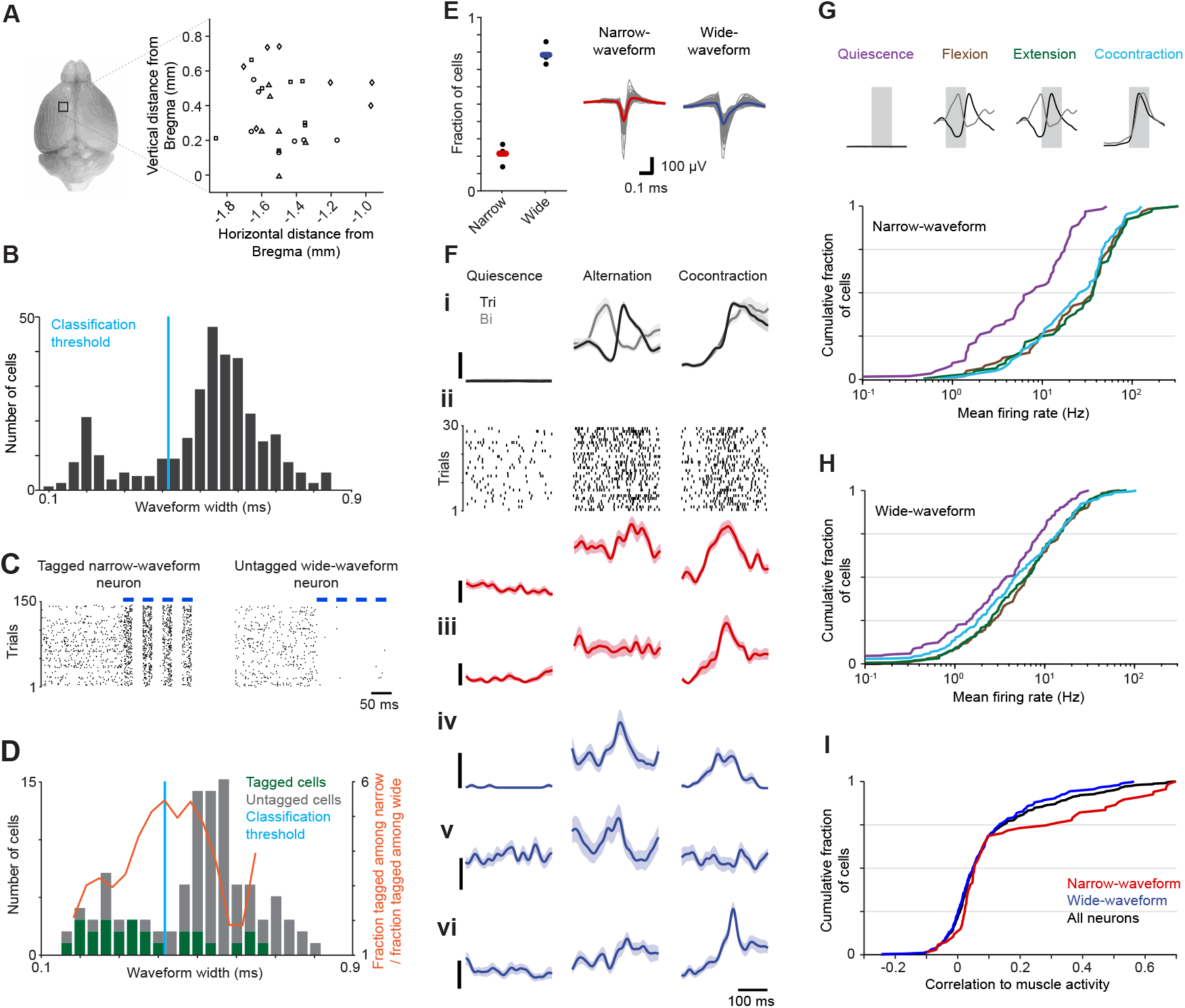
Neural activity during task performance. (A) Region of left motor cortex from which probe recordings were made (left) and recording sites for each mouse (right; circle, diamond, square, and triangle symbols signify each of 4 mice). (B) Distribution of trough to peak waveform widths for neurons recorded from cortical sites shown in (A). (C) Spike rasters from an optically tagged narrow-waveform neuron (left) and an untagged wide-waveform neuron (right) during light stimulation (blue bars) in VGAT-ChR2-EYFP mice. (D) Distribution of waveform widths of neurons that were optically tagged (26/104, green) or untagged (78/104, gray). The waveform width classification threshold (blue) was identified as 0.42 ms. (E) Fractions of neurons assigned to each subtype for each mouse (left; black dots). Colored bars show means across mice. Mean waveforms (right) for narrow-(n = 67) and wide-waveform neurons (n = 249), with overall mean bold and colored. (F) Trial-averaged EMG and neural activity ± SEM (n = 30 trials) during muscle quiescence (left column), alternation (middle), and cocontraction (right) epochs. (i) EMG activity in triceps (Tri, black) and biceps (Bi, gray). (ii) Spike rasters of a narrow-waveform neuron and trial-averaged firing rates for that neuron immediately below. (iii-vi) Trial-averaged firing rates for an additional narrow-waveform (iii, red) and three wide-waveform neurons (iv-vi, blue). Scale bars represent 10% of maximal EMG activity (top) and 20 Hz (below), respectively. (G-H) Cumulative histograms of mean firing rates during quiescence, flexion, extension, and cocontraction for narrow-(G) and wide-(H) waveform neurons. Means are measured as the mean of the trial-averaged time series segment overlaid with gray (top). Histograms in (G-I) exclude with firing rates that did not rise above 1 Hz in any epoch. (I) Cumulative histogram of the correlation of neuronal firing rates with muscle activity across the entire session for narrow-waveform (red), wide-waveform (blue), and all neurons (black).

To separately assess the activity of putative excitatory and inhibitory neurons from our recordings, we used the relationship between trough-to-peak waveform width and neuronal identity to identify narrow-waveform, putative inhibitory interneurons and wide-waveform, putative pyramidal neurons (Barthó et al., 2004; McCormick et al., 1985; Figure 2B). To guide classification, we recorded activity in L5b of CFA during exposure to pulsed 473 nm light in a separate cohort of mice expressing channelrhodopsin2 in vGAT^on^ cortical inhibitory neurons (Zhao et al., 2011). Neurons were designated “tagged” and thus inhibitory if the latency to spike after light onset was significantly reduced compared to equivalent periods without light (Kvitsiani et al., 2013; Figure 2C). We then identified a waveform width for separating putative inhibitory and pyramidal neurons that maximized the ratio between the fraction of tagged neurons among the narrow-waveform population and the fraction of tagged neurons among the wide-waveform population (Figure 2D). Using this assignment method, 22% ± 3% of neurons recorded during behavioral performance were narrow-waveform and 78% ± 3% of neurons were wide-waveform (Figure 2E), consistent with previous observations (Beaulieu, 1993; Guo et al., 2014a; Miri et al., 2017).

The activity of narrow- and wide-waveform neurons agreed with that seen previously in primary motor cortical regions during limb movement in mice and other mammals. To compute average responses across multiple instances of the same behavior, we used triceps and biceps activity to identify 30 similar time series segments (“trials”) per task epoch. Only neurons that did not exhibit a substantial difference in mean firing rate between the two alternation epochs were included in this and subsequent analyses (Figure S2), but we note that omitting these cells did not affect the essence of our results. Trial-averaged neuronal firing rates exhibited a broad diversity of patterns (Figure 2F; Churchland and Shenoy, 2007). Neural responses often had substantial correlation with muscle activity (Kargo and Nitz, 2004; e.g., compare the neuron in 2Fiv with triceps activity), but correlations could be prominent for one task and not the other (e.g., the neuron in 2Fv). On average, narrow-waveform neurons had higher firing rates and showed a greater difference between mean firing rates during muscle quiescence (mean ± SEM, 10.5 ± 1.3 spikes/s) and task engagement (Alt: 39.1 ± 5.3, Cc: 27.1 ± 3.1 spikes/s, n=67 neurons) relative to wide-waveform neurons (Qui: 6.0 ± 0.4, Alt: 10.0 ± 0.7, Cc: 8.9 ± 0.7 spikes/s, n=249 neurons; Figure 2G-H; Isomura et al., 2009; Kaufman et al., 2013). Activity in both neuronal subsets was higher on average during task performance compared to periods of muscle quiescence (Figure 2G-I; Estebanez et al., 2017; Isomura et al., 2009; Miri et al., 2017).

### Motor cortical activity does not conform to existing hypotheses

Previously studies have described distinct subpopulations of motor cortical neurons that modulate their activity in relation to cocontraction but not flexion or extension (Fetz and Cheney, 1987; Humphrey and Reed, 1983). If cells do form distinct, task-specific subpopulations, plots of mean firing rates during alternation versus those during cocontraction would show clusters of points displaced in either direction from the identity line. Here and in the analysis that follows, we focused on neurons that (a) had mean firing rate > 1 Hz during either alternation or cocontraction, (b) were consistently task-related (firing rate change between the two alternation epochs < 50%), and (c) were stably recorded (absolute slope of the Euclidean distance between waveform at beginning and end of session < 0.2), yielding 325 of 804 neurons from 4 mice (Methods: Firing Rate Calculation, Table 1). Forgoing these three criteria had no meaningful effect on our results.

**Table 1:** Elbow trials

| Mouse ID | # recorded units | # units after behavioral session subselection | # units after retaining $\geq 1$ Hz firing rate in at least 1 epoch | # units after filtering for $>1.5$ fold Alt1-Alt2 FR difference | # units after filtering for high waveform pdist (probe instability) |
| --- | --- | --- | --- | --- | --- |
| A32 | 320 | 240 | 224 | 90 | 86 |
| A33 | 589 | 205 | 196 | 93 | 92 |
| A35 | 444 | 215 | 202 | 72 | 71 |
| A36 | 233 | 144 | 139 | 70 | 67 |
| Total | 1586 | 804 | 761 | 325 | 316 |

**Table 2:** Wrist trials

| Mouse ID | # recorded units | # units after behavioral session subselection | # units after retaining $\geq 1$ Hz firing rate in at least 1 epoch | # units after filtering for $>1.5$ fold Alt1-Alt2 FR difference | # units after filtering for high waveform pdist (probe instability) |
| --- | --- | --- | --- | --- | --- |
| A32 |  |  | 173 | 86 | 81 |
| A33 |  |  | 156 | 68 | 67 |
| A35 |  |  | 158 | 64 | 63 |
| Total |  |  | 487 | 218 | 211 |

We observed no clusters either when considering the wide-waveform subpopulation, which includes putative motor cortical output neurons (Figure 3A), or all neurons (Figure 3B). Nor was evidence of clusters apparent in histograms of the distances of points from the identity line, and the distribution of these distances was not significantly multimodal (Hartigan’s Dip test, dip = 0.012, p = 1 (wide); dip = 0.011, p = 1 (all); Figure 3C-D). Moreover, measurement of the firing rate in one task relative to the overall firing rate for each neuron (specificity index) demonstrates a continuum of task specificity for alternation and cocontraction among wide-waveform neurons (Figure 3E) and all neurons (Figure 3F). These results indicate that task-specific neurons are not a prominent feature of motor cortical L5b.

**Figure 3.**
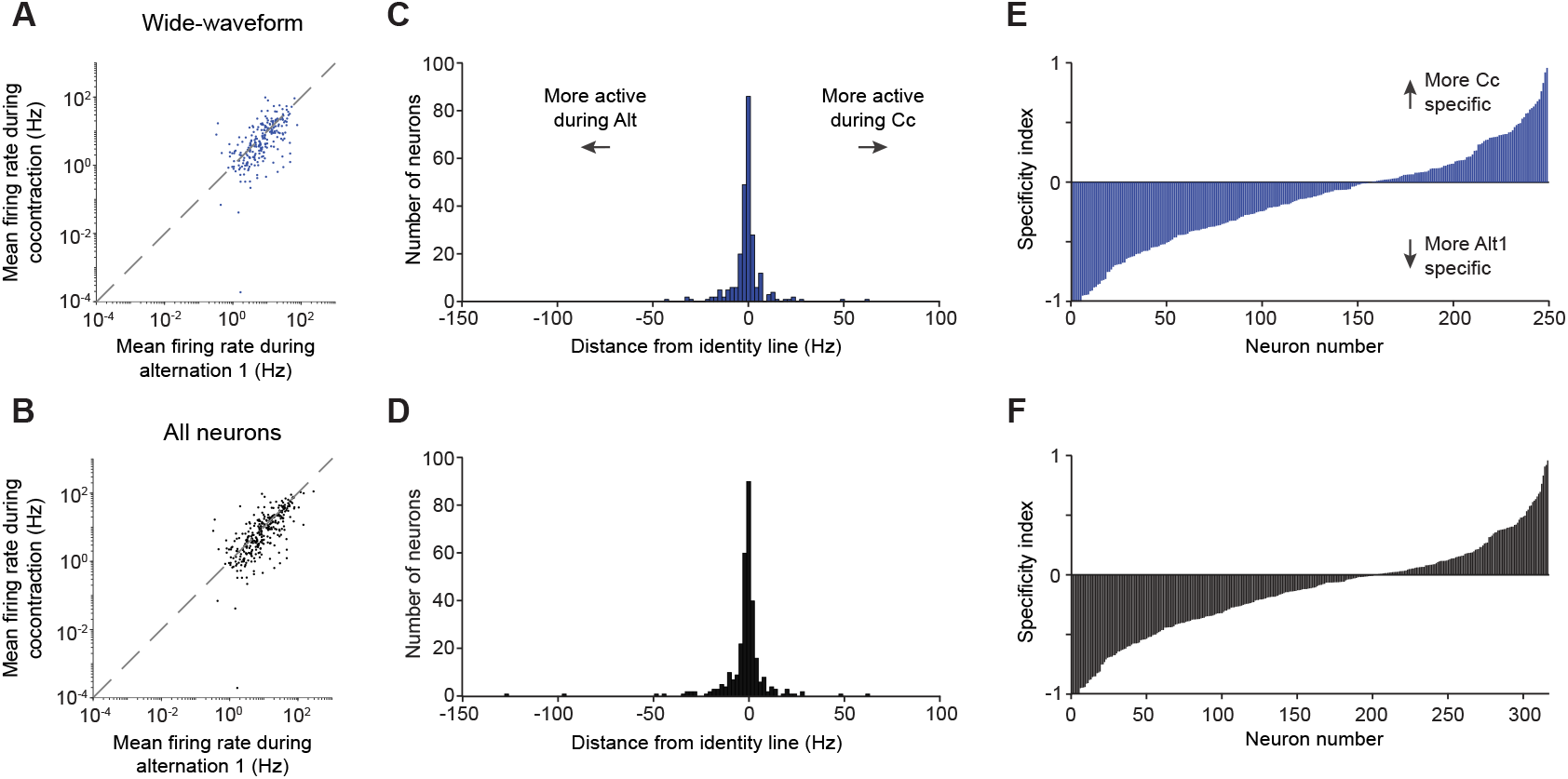
Neurons demonstrate a continuum of task specificity. (A-B) Mean firing rate during the first epoch of alternation versus mean firing rate during cocontraction for wide-waveform neurons (A) and all neurons (B). (C-D) Distribution of the distance of each point in (A) and (B) from the identity line. (E-F) Specificity index for wide-waveform (E) and all neurons (F), calculated by taking the difference between the mean firing rates during two tasks divided by their sum. Neurons are separately ordered by ascending index value in each plot.

The idea that motor cortex mediates different muscle activity patterns through activation of flexion- and extension-related populations implies a consistent relationship between motor cortical output and muscle activity during alternation and cocontraction. To test this, we used ridge regression to fit trial-averaged muscle activity with a linear sum of trial-averaged neural activity. We first fit each muscle’s activity using wide-waveform neuron activity during the first epoch of the alternation task (Alt1), then evaluated performance during three situations: held-out data for Alt1, the cocontraction epoch (Cc), and the second alternation epoch (Alt2). We found these models generalized well to activity from alternation (R^2^ ± SEM = 0.39 ± 0.08 for held-out Alt1, 0.35 ± 0.08 for Alt2, n = 4 mice). In contrast, there was essentially no generalization to Cc (0.002 ± 0.02; Figure 4A-B). This effect – better generalization during alternation versus cocontraction – varied in magnitude across all six recorded muscles but was always present (Figure S3C). The same effect was seen in reverse when models were trained using Cc data: models performed well on held-out Cc data (0.75 ± 0.04) but poorly fit data from both Alt1 (−0.82 ± 0.33) and Alt2 (−1.08 ± 0.44; Figure 4C-D). The low R^2^ values below 0 for fits to alternation epochs indicate that the models fit worse than a horizontal line, a clear sign that models trained to cocontraction epochs are inappropriate for alternation epochs. Similar results were seen if models employed all neurons, rather than only wide-waveform neurons (Figure 4B,D).

**Figure 4.**
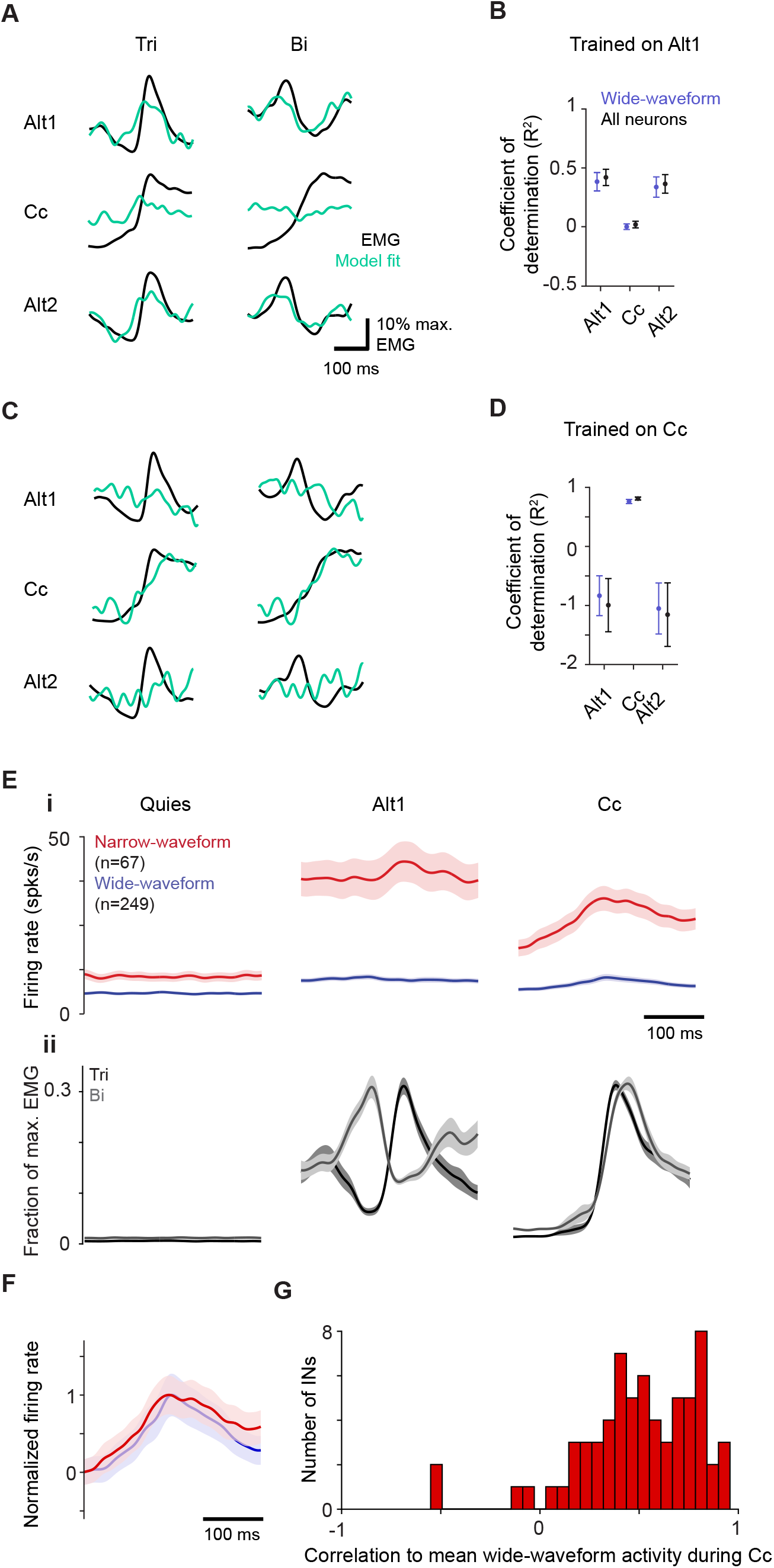
Neural activity does not conform to existing hypotheses. (A, C) Example of ridge regression model fit to trial-averaged triceps (Tri) and biceps (Bi) activity for models trained on activity from the first alternation epoch (A) or the cocontraction epoch (C). (B, D) Mean ± SEM (n = 4 mice) coefficient of determination for models trained to fit activity for all six muscles from the first alternation epoch (B) and cocontraction (D) using wide-waveform or all neurons when tested on activity from the first alternation epoch (Alt1), cocontraction (Cc), and the second alternation epoch (Alt2). (E) Trial-average activity ± SEM during muscle quiescence, alternation, and cocontraction of narrow- (i; n = 67 neurons from 4 mice) and of wide-waveform neurons (ii; n = 249 neurons). (F) Trial-average activity ± SEM of narrow- (red) and wide- (blue) waveform neurons normalized to their respective minima and maxima averaged across all 4 mice. (G) Distribution of correlation coefficients for each narrow-waveform neuron’s activity to the mean wide-waveform activity during cocontraction (n = 67 neurons from 4 mice).

We then considered whether an adequate model for both epoch types could be found by training models on data from both epochs simultaneously. However, models trained on both the Alt1 and Cc activity performed well for Cc (0.71 ± 0.06) but performed very poorly for alternation (Alt1 −0.10 ± 0.27, Alt2 −0.20 ± 0.22; Figure S3A). Since the model could be fitting to cocontraction well at the expense of alternation, we further sought a good fit to data from both epochs by encouraging a better fit to alternation data, artificially reducing the scale of neural and muscle activity during cocontraction. However, a good fit for both epochs was never obtained (Figure S3B). Taken together, these regression results indicate that the relationship between motor cortical and muscle activity varies between alternation and cocontraction, arguing against the existence of populations consistently related to flexion or extension.

Other studies arguing for the existence of consistent flexor- and extensor-related populations propose that their coactivation results from a reduction in intracortical reciprocal inhibition between them (Capaday, 2004; Ethier et al., 2007; Matsumura et al., 1991). This view predicts that during cocontraction, the activity of interneurons and pyramidal neurons will be negatively correlated, and overall pyramidal neuron activity will increase: interneuron activity decreases, allowing both flexion- and extension-related pyramidal populations to simultaneously increase their activity. We did find that the activity of narrow-waveform neurons was overall lower during cocontraction versus alternation (75.8% of mean alternation firing rate; Figure 4Ei). However, rather than decreasing, narrow-waveform activity increased at cocontraction onset. Moreover, the total activity of wide-waveform neurons did not increase during cocontraction (98.4% of mean alternation firing rate; Figure 4Eii). This absence of an increase during cocontraction also argues against the possibility that interneurons in other layers mediate the hypothesized intracortical reciprocal inhibition between flexor- and extensor-related populations. Rather than a negative correlation, a positive correlation is readily apparent when comparing the normalized mean firing rates between the narrow- and wide-waveform populations (Pearson correlation = 0.97; Figure 4F). In addition, very few narrow-waveform neurons had activity that was anticorrelated with average wide-waveform activity (Figure 4G). Only 4/67 narrow-waveform neurons had negative correlations, among a population with a mean ± SEM correlation of 0.50 ± 0.04. Together, these results suggest that in motor cortex a relief of intracortical inhibition does not occur during cocontraction, but instead excitation and inhibition remain balanced.

### Changes in the covariation of motor cortical activity across tasks

Another possible mechanism for the mediation of different muscle activation patterns by motor cortex is a change in the covariation of its output activity patterns. Variation along certain directions in neural activity space could differentially drive changes in spinal circuits to facilitate either cocontraction or alternation. This mechanism would also explain the finding that linear models generalize poorly across tasks even though descending connections presumably remain fixed. To test this mechanism, we used principal component analysis to identify activity space directions along which motor cortical activity varies strongly during alternation and cocontraction. The first several principal components of trial-averaged firing rates define directions that account for a large fraction of activity variation across the population (covariation). The first six principal components during each task epoch accounted for approximately 90% of the firing rate variance (mean ± SEM, Alt1 variance capture = 89.60% ± 0.01%, Cc variance capture = 92.56% ± 0.02%, Alt2 variance capture = 89.75% ± 0.02%; Figure 5A).

**Figure 5.**
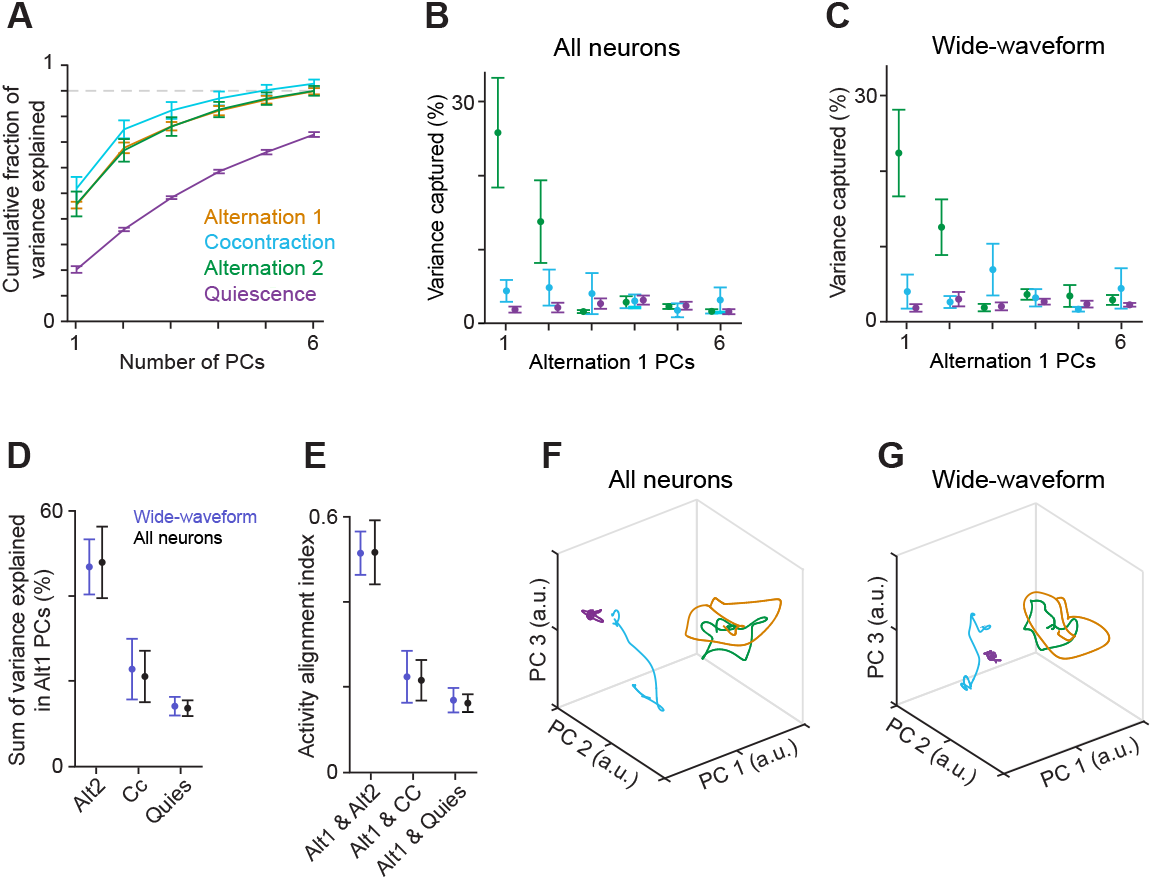
Task-specific motor cortical activity covariation. (A-C) Mean ± SEM (n = 4 mice) cumulative variance explained by the top principal components (PCs) for neural activity from alternation 1, cocontraction, alternation 2, and muscle quiescence (A) and variance explained by the top PCs of alternation 1 among all neurons (B) and wide-waveform neurons (C). (D-E) For wide-waveform and all neurons, mean ± SEM (n = 4 mice) of the sum of variance explained by the top 6 alternation 1 PCs (D) and of the alignment of firing rates in alternation 1 with other epochs (E). (F-G) Neural activity projected into the top three PCs of the shared alternation 1 and cocontraction activity space for all (F) and wide-waveform neurons (G). Colors consistent with A-C.

Consistent with the above mechanism, the neural activity subspaces inhabited during alternation and cocontraction were substantially different. The Alt1 principal components accounted for a considerable amount of the firing rate covariation during Alt2 (Figure 5B, green), but little of the covariation during cocontraction (blue) or quiescence (purple). This was true for both all neurons and the wide-waveform subset (Figure 5C). This effect was particularly pronounced for the top two principal components but was also apparent for the top six principal components collectively (Figure 5D). The variance captured during cocontraction was far less than that captured during Alt2, and only slightly more than that captured during quiescence, which provides an estimate of baseline variance capture. To control for any differences in the variance captured by top principal components across tasks, we also computed the subspace ‘alignment index,’ the ratio between the variance in a given epoch accounted for by components computed for a second epoch and that accounted for by components computed for the given epoch (Figure 5E; Elsayed et al., 2016). For all neurons, the alignment index for both alternation epochs was 0.52 ± 0.07 (mean ± SEM), while this index for Alt1 and Cc (0.22 ± 0.05) was only slightly higher than that for Alt1 and muscle quiescence (0.16 ± 0.02). Thus, the ability of the alternation subspace to capture neural activity during cocontraction was not much better than its ability to capture neural activity in the absence of muscle output.

Further evidence for changes in motor cortical activity covariation across tasks was found when we calculated principal components for activity in both Alt1 and Cc epochs collectively. Projection of activity onto these principal components illustrated that distinct components captured the bulk of activity variance during each of the two tasks (Figure 5F-G). In general, two components captured much of the cyclic activity variation during alternation, while a separate third component captured much more of the variance during cocontraction.

Given that motor cortical and muscle activity are correlated, we checked whether changes in neural activity covariation simply reflect the change from negatively correlated flexor-extensor muscle activity during alternation to positively correlated activity during cocontraction. To compare changes in covariation between alternation and cocontraction for motor cortical and muscle activity, we first equalized the dimensionality of both activities by projecting them onto the top six principal components computed collectively for Alt1 and Cc. Thus, both neural and muscle activity were exactly six-dimensional, and so were guaranteed to have a unity alignment index when considering six-dimensional subspaces. We then computed the alignment index for this projected activity during Alt1 and Cc using varying numbers of components, finding that the alignment index was consistently lower for the neural data (Figure S4A,B). This indicates that the observed changes in motor cortical activity covariation cannot be explained simply by changes in muscle activity covariation alone, and more generally that the population-level structure of motor cortical activity can be very different from that of muscles (Russo et al., 2018).

We also verified that changes in neural activity covariation did not reflect changes only in activity unrelated to muscle activity. Although the subspace alignment for Alt1 and Cc was similar to the alignment for either epoch and quiescence, the subspace shared between alternation and cocontraction could be responsible for controlling muscles. Changes in activity subspace might then reflect, for example, cognitive processes not directly related to muscle control and the differences in sensory context between tasks. These latter differences include a greater degree of limb movement during the alternation task, the higher rung location during the cocontraction task, and different static visual cues that signal which task is expected. To examine only muscle-correlated motor cortical activity, we projected trial-averaged activity onto a six-dimensional subspace defined by the ridge regression fits to activity of each of the six recorded limb muscles. The same change in subspace occupancy reflected in variance capture by top principal components and alignment index was observed for activity after this projection (Figure S4C,D). This was true both for all neurons and the wide-waveform subset. This indicates that activity subspaces inhabited during alternation and cocontraction are substantially different specifically for muscle-related motor cortical activity.

### Similar motor cortical activity structure during wrist muscle alternation and cocontraction

To see how well the findings described above hold for the cortical control of other joints, we then sought to replicate these findings for alternation and cocontraction of wrist antagonists (extensor digitorum communis, EDC; and palmaris longus, PL). In the three of four mice used for neural recording that expressed cocontraction at the wrist in our paradigm, trial sets were identified for quiescence, alternation, and cocontraction from each recording session (Figure 6A).

**Figure 6.**
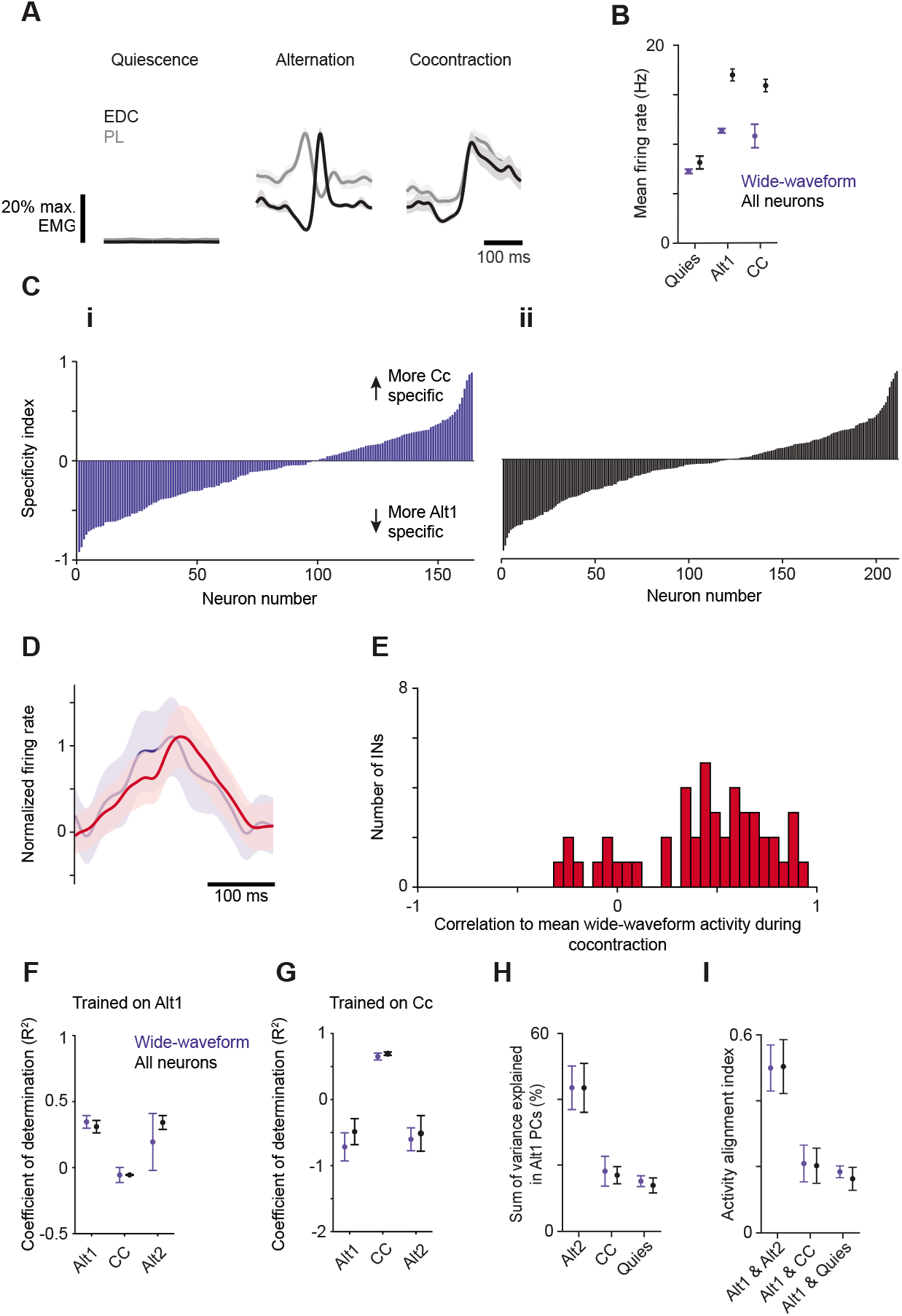
Motor cortical activity during wrist alternation and cocontraction also reflects task-specific covariation. (A) Trial-averaged muscle activity ± SEM (n = 30 trials) for EDC and PL during muscle quiescence (left), alternation (middle), and cocontraction (right) epochs. (B) Mean ± SEM firing rates from 3 animals during three task epochs for wide-waveform (n = 164 neurons) and all neurons (n = 211 neurons). (C) Specificity index for neurons separately in ascending order for wide-waveform (i) and all neurons (ii). (D) Trial-averaged activity ± SEM of narrow- (red; n = 47 neurons) and wide- (blue; n = 164 neurons) waveform neurons normalized to their respective minima and maxima. (E) Distribution of correlation coefficients for each narrow-waveform neuron’s activity to the mean wide-waveform activity during cocontraction. (F, G) Mean ± SEM (n = 3 mice) coefficient of determination (R^2^) for models trained to fit activity for all six muscles from the first alternation epoch (F) and cocontraction (G) using wide-waveform or all neurons when tested on activity from the first alternation epoch (Alt1), cocontraction (Cc), and the second alternation epoch (Alt2). (H, I) For all (black) and wide-waveform neurons (blue), mean ± SEM (n = 3 mice) of the sum of variance explained by the top 6 alternation 1 principal components (PCs; H) and of the alignment of firing rates in alternation 1 with other epochs (I).

Consistent with findings at the elbow, mean firing rates during alternation and cocontraction exceeded that during quiescence for both wide-waveform and all neurons (Figure 6B). Specificity indices demonstrated a continuum of task specificity across neurons rather than any apparent clusters of task-specific neurons (Figure 6Ci, ii). The mean normalized firing rates of wide- and narrow-waveform neurons were highly correlated (Figure 6D), and on a per neuron basis, activity in the vast majority of narrow-waveform neurons was correlated with mean wide-waveform activity (Figure 6E).

For the wrist, we again found that alternation and cocontraction do not rely on activation of the same flexion- and extension-related populations. We applied ridge regression to fit trial-averaged muscle activity for alternation and cocontraction tasks with linear sums of trial-averaged neural activity. We found that a model trained on Alt1 data generalized to held-out Alt1 and Alt2 activity (R2 ± SEM = 0.310 ± 0.047 for held-out Alt1, 0.341 ± Y for Alt2 in all neurons; 0.346 ± 0.477 for held-out Alt1, 0.194 ± 0.215 for Alt2 in wide-waveform neurons; n = 3 mice) but failed on Cc activity (in all neurons −0.055 ± 0.003; in wide-waveform neurons −0.056 ± 0.057; Figure 6F). Conversely, a model trained on Cc activity performed well on held-out Cc trials (in all neurons, 0.685 ± 0.0255; in wide-waveform neurons 0.642 ± 0.522; Figure 6G) but failed to generalize to alternation (in all neurons, −0.492 ± 0.196 for held-out Alt1, −0.518 ± 0.269 for Alt2; in wide-waveform neurons, −0.7218 ± 0.212 for held-out Alt1, −0.607 ± 0.1729 for Alt2).

We again found dramatic changes in the covariation of motor cortical activity patterns between alternation and cocontraction. We found that the sum of variance explained in the top 6 Alt1 principal components accounted for substantial variance in Alt2 (all neurons, 0.434 ± 0.074; wide-waveform neurons, 0.434 ± 0.066) but little of that in cocontraction (all neurons, 0.170 ± 0.026; wide-waveform neurons, 0.182 ± 0.0449; Figure 6H). The subspace alignment index was much higher between alternation epochs (all neurons, 0.504 ± 0.081; wide-waveform neurons, 0.450 ± 0.070) than between alternation and cocontraction (all neurons, 0.204 ± 0.053; wide-waveform neurons, 0.211 ± 0.055). These latter alignment indices in turn were only slightly higher than those for alternation and muscle quiescence (all neurons, 0.165 ± 0.035; wide-waveform neurons, 0.186 ± 0.018; Figure 6I). Collectively, these results indicate that similar neural dynamics underlie motor cortical control of elbow and wrist muscles.

### Changes in corticospinal activity covariation across tasks

We then addressed whether findings for L5b putative pyramidal neurons held specifically for a primary contributor to motor cortical output: corticospinal neurons. To induce expression of the genetically encoded calcium indicator GCaMP6f specifically in this neuronal population, an rAAV2-retro was injected into cervical spinal segments (Figure 7A). As mice performed in our behavioral paradigm, two-photon imaging was used to measure fluorescence in cross-sections of apical trunk dendrites in horizontal planes ∼350 µm below pia (Figure 7B-C; Mittmann et al., 2011; Peters et al., 2017). Calcium fluctuations and spiking were estimated for identified cross-sections (Figure 7D; Giovannucci et al., 2019; Pnevmatikakis and Giovannucci, 2017). Sets of similar behavioral segments were identified from simultaneous EMG recordings during each task epoch, and trial-averaged estimated spiking was computed for each dendritic section whose activity met selection criteria (n = 194 across 9 imaging windows in 2 mice; Figure S5A-E; see Methods). Detected spike rates were lower for fluorescence measurements from corticospinal neurons compared with electrical recording of wide-waveform neurons, likely due at least in part to limited indicator sensitivity relative to the size of spike-induced free calcium increases in indicator-loaded cells, coupled with the sparse firing of many pyramidal neurons.

**Figure 7.**
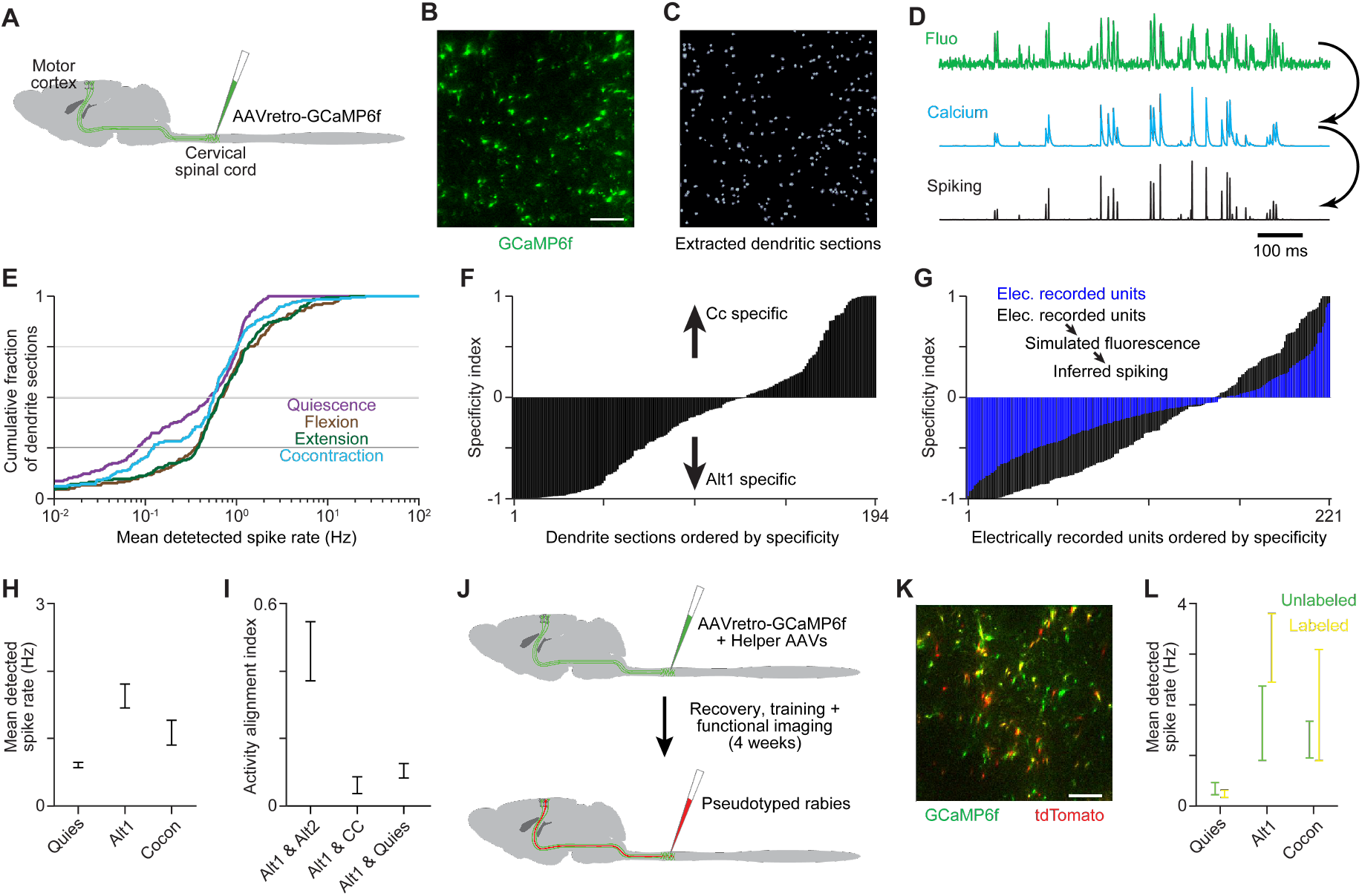
Corticospinal neuron activity during alternation and cocontraction. (A) An rAAV2-retro was injected into cervical spinal segments to induce GCaMP6f expression in corticospinal neurons. (B-C) Example of fluorescence (B) and algorithmically extracted dendritic cross-sections (C) from a horizontal imaging region ∼350 µm below pia. Scale bar = 50 µm. (D) Example fluorescence, estimated calcium change, and estimated spiking time series. (E) Cumulative histogram of detected spike rates for qualifying dendritic sections (n = 194) during distinct behavioral epochs. (F) Specificity index for qualifying dendritic sections, in ascending order. (G) Specificity index for electrically recorded putative pyramidal neurons, and the same neurons after simulating calcium-sensitive fluorescence based on their spiking and inferring spiking from simulated fluorescence. (H) Mean ± SEM detected spike rate across qualifying dendritic sections during distinct behavioral epochs. (I) Mean ± SEM (n = 9 imaging planes) activity alignment for corticospinal dendritic sections. (J) Schematic of injections for imaging experiments in which rabies-driven tdTomato labeling of GAD+ spinal neuron-contacting corticospinal neurons was registered to GCaMP6f after functional measurements were completed. (K) Example of GCaMPf and tdTomato fluorescence from a horizontal imaging region. Scale bar = 50 µm. (L) Mean ± SEM for tdTomato labeled (n = 28) and unlabeled (n = 19) dendritic sections during quiescence, alternation, and cocontraction. Neither group showed significant changes between alternation and cocontraction (two-tailed t-test, p > 0.05).

Findings for electrically recorded wide-waveform neurons largely held for corticospinal neurons. First, distributions of mean activity across dendritic sections showed less activity when muscles were inactive than during alternation or cocontraction (Figure 7E). Second, a specificity index comparing activity modulation during alternation and cocontraction again exhibited a continuum of values (Figure 7F). We did note that extreme index values were more prevalent than for electrically recorded neurons, but we found that this difference could be due to limited indicator sensitivity and aspects of the relationship between actual and inferred spiking that are not well captured by existing spike inference methods (Greenberg et al., 2018). We showed this by simulating calcium-sensitive fluorescence and inferred spiking measurements for our electrically recorded wide-waveform neurons and recalculating their specificity indices (Figure 7G). Importantly, simulation parameters were tuned to match the rate of spike detection and not the distribution of specificity indices itself.

Third, we found that aggregate corticospinal neuron spike rates were not higher during cocontraction compared to alternation (Figure 7H), arguing against the coactivation of consistent flexion- and extension-related output populations. Lastly and most strikingly, the average alignment between activity subspaces in At1 and Cc was even lower for corticospinal neurons than for the electrically recorded L5b population. Subspace alignment for corticospinal neurons was much lower for Alt1 and Cc compared with Alt1 and Alt2 and was on par with the alignment between Alt1 and quiescence (Figure 7I). Thus, corticospinal neurons appear to influence alternation and cocontraction primarily through changes in activity covariation.

Since previous evidence has implicated increased presynaptic inhibition of sensory afferent terminals in facilitating cocontraction (Nielsen et al., 1993), we examined whether corticospinal neurons that contact spinal interneurons mediating presynaptic inhibition are preferentially activated during cocontraction. We used a transsynaptic rabies-based approach to identify a subset of GCaMP6f-labeled corticospinal neurons that contact GAD2+ neurons in cervical spinal cord (Figure 7J,K), a population enriched for presynaptic inhibitory interneurons (Fink et al., 2014). We found that the mean activity of GAD2+ interneuron-contacting corticospinal neurons was not significantly increased during cocontraction compared to alternation (Figure 7L). This finding is consistent with a motor cortical mediation of cocontraction that relies not on a specific population of output neurons, but instead on a change in activity covariation across output neurons.

## DISCUSSION

We have examined how motor cortex mediates tasks comprising different muscle activation patterns using a novel motor cortex-dependent paradigm that elicits forelimb flexor-extensor alternation and cocontraction in mice. Critical here was our use of real-time detection of muscle activity features to rapidly trigger reward, which presents a new avenue for training mice to generate specific motor outputs. Population-level analyses of activity electrically recorded in L5b and optically recorded from corticospinal neurons did not support a view grounded in a muscle-based cellular organization that motor cortical influence relies on similar flexion- and extension-related populations during both alternation and cocontraction. Our analysis also failed to support a view grounded in a task-based cellular organization that these forms of muscle activation rely on dedicated neuronal populations.

Instead, our results support the emerging hypothesis that tasks requiring different muscle activation patterns are mediated through task-specific neural activity covariation (Figure 1C), which we observed for all recorded L5b neurons and specifically for wide-waveform putative pyramidal and corticospinal subsets. In this view, the strong variation along certain activity space directions seen only during cocontraction is what promotes a reduction in reciprocal inhibition and enables coactivation of antagonist motor pools. Spinal interneurons downstream of motor cortical output would respond preferentially to such activity covariation due to the weights of intervening synapses, leading to appropriate modulation of spinal interneurons and motor neurons. Our findings support the view that motor cortical control relies on activity coordinated at the population level (Churchland et al., 2012; Gallego et al., 2017; Sussillo et al., 2015), such that distinct covariation in motor cortical output drives distinct functions (Druckmann and Chklovskii, 2012; Miri et al., 2017).

### Contrast with existing hypotheses

Our results do not support a model for the control of voluntary cocontraction based on the activity of a task-specific subpopulation. If such a subpopulation existed, its activity should be differentiable from the activity of the overall population. However, our population-level analysis did not detect such a subset, instead revealing a continuum of task-specificity. A minority of neurons predominantly or exclusively active during cocontraction, observed here and by others (Fetz and Cheney, 1987; Humphrey and Reed, 1983), appear to exist at the extremes of this continuum.

Our findings also failed to support the idea that voluntary cocontraction is produced by an attenuation of intracortical inhibition in motor cortex that permits coactivation of distinct pyramidal neuron subsets dedicated to the activation of either flexor or extensor muscles (Capaday et al., 1998; Ethier et al., 2007; Matsumura et al., 1991; Schneider et al., 2002). Were this true, best-fit models of motor cortical activity to muscle activity would generalize between alternation and cocontraction. Instead, we found models did not generalize, and those trained on data from both tasks did a comparatively poor job of fitting muscle activity. The change in the relationship between motor cortical and muscle activities appears general and not particular to best-fit models, as flexor-extensor alternation and cocontraction were associated with activity in motor cortex along distinct directions in neural activity space. Thus, any model trained on one task is likely guaranteed to generalize poorly.

Also contradicting the above hypothesis was our observation that the firing rates of putative inhibitory interneurons increased in parallel with putative pyramidal activity, which itself was not higher during the cocontraction task compared to the alternation task. This latter finding also argues against the possibility that interneurons in other layers may be mediating intracortical inhibition between flexor- and extensor-related pyramidal neuron populations in L5b, the reduction of which would allow both pyramidal populations to coactivate (Figure 1B). This would lead to an overall increase in pyramidal cell activity during cocontraction, which was not observed. The balance of cortical excitation and inhibition we saw also agrees with recent observations of increases in the activity of motor cortical interneurons at movement onset in monkey, rat, and mouse (Estebanez et al., 2017; Isomura et al., 2009; Kaufman et al., 2013).

### Task-specific activity covariation in motor cortex

Our results demonstrating a profound change in motor cortical activity covariation between the alternation and cocontraction tasks suggest a new model for motor cortical influence on these muscle activation patterns. In this model, cocontraction-specific activity covariation in motor cortical output promotes the reduction in reciprocal inhibition and coactivation of antagonist motor pools in the spinal cord (Figure 1C). The magnitude of the observed change in covariation and the likelihood that output neurons are not under-represented in recorded populations (Miri et al., 2017) suggest that covariation changes hold for motor cortical output neurons. Moreover, here we specifically demonstrate a profound covariation change in a key motor cortical output population, the corticospinal neurons. Our finding that the majority of corticospinal neurons are more active during muscle contraction also contrasts with the finding that a majority of corticospinal neurons decrease their activity at movement onset (Peters et al., 2017), a discrepancy that could be explained by muscle activity preceding movement cues.

The change in activity covariation we observed appears to be a robust feature of motor cortical activity across the present tasks. Given the relatively low dimensionality of activity during either task, if activity covariance was constant between tasks so that any observed changes reflect chance variation, a much smaller covariance change would be expected (Sadtler et al., 2014). Indeed, we found the covariance change between two epochs of alternation to be much lower. We have also shown that the covariance change between the alternation and cocontraction tasks is not simply a consequence of the change in muscle covariation between these tasks (Figure S4A,B).

Our findings raise the question of what it is that causes the observed changes in activity covariation. Given the strong interconnections between primary motor and somatosensory cortices and the relevance of sensory feedback for generating motor commands, it is possible that sensory feedback, or other inputs to motor cortex that differ between task contexts, contribute to the differences in neural variation observed. However, it is unlikely that our observations reflect a sensory representation unrelated to muscle control. As argued previously (Miri et al., 2017), the degree of difference between activity subspaces is large enough that any substantial component of L5b motor cortical activity, such as the components carried by pyramidal tract and intratelencephalic neurons that influence muscle activity, should differ as well. The activity subspace overlap we observe is similar to that seen between task performance and muscle quiescence, which can be viewed as a lower bound on the degree of overlap between two task contexts. Most importantly, we extend our findings here to corticospinal neurons, where subspace overlap was again on par with that seen between task performance and muscle quiescence.

That task-specificity in motor cortical influence relies on population-wide changes in activity covariation contrasts with historical views of motor cortical organization in which functions, be it the control of particular movement types or sets of muscles, were attributed to distinct cellular subpopulations (Graziano et al., 2006; Penfield and Rasmussen, 1950). More recent studies using calcium indicator imaging in mice have also suggested the existence of task-specific neuronal subpopulations (Dombeck et al., 2009; Komiyama et al., 2010). We note that the nonlinear encoding of activity by calcium indicators could exaggerate the appearance of categorical distinctions between cells. Our own results both with electrical recording and imaging show a similar degree of task-specificity across neurons, one which coexists with strikingly different patterns of activity covariation. We suspect that our finding of a continuum of task-specificity implies that neurons that appear task-selective exist on the margins of this continuum.

### Task-specific covariation more broadly

Our findings add to a number of recent studies demonstrating that different tasks are associated with distinct motor cortical activity covariation. Covariation changes may underlie how motor cortical output differentially engages downstream circuits during tasks with different motor cortical involvement (Miri et al., 2017). Differences in motor cortical activity covariation are also observed between forward and backward arm cycling (Russo et al., 2018), movement preparation and subsequent execution (Elsayed et al., 2016; Kaufman et al., 2014), movement of the ipsilateral and contralateral limbs (Ames and Churchland, 2019), and different tasks performed via brain-machine interface (Schroeder et al., 2021). These differences in activity covariation may help ensure independence between different cortical functions (Mante et al., 2013; Pandarinath et al., 2018; Perich et al., 2018). In the case of alternation and cocontraction, distinct subcortical circuits controlling limb movement and steadiness (Shadmehr, 2017; Albert et al., 2019; Saliba et al., 2020) could be independently controlled by distinct motor cortical output covariation. It should also be noted that many movements involve simultaneous alternation and cocontraction (Tilney & Pike, 1925), and it has been suggested that the separate modes that control each are activated in parallel (Feldman, 1980; Scheidt & Ghez, 2007; Shadmehr, 2017).

Task-specific activity covariation has been found in neuronal populations beyond the motor cortex. This includes findings of task-specific cortical activity dynamics during decision-making (Pinto et al., 2019), task-specific representation of stimulus and choice in parietal cortex (Pho et al., 2018), and changes in the encoding of eye position in the oculomotor neural integrator across different types of eye movement (Daie et al., 2015). Such covariation changes also resemble those observed across hippocampal place cells as the surrounding environment changes (Leutgeb et al., 2005). Different patterns of activity covariation defining what have been termed “communication subspaces” appear to underlie the selective routing of signals from a given cortical area to particular targets (Semedo et al., 2019; Perich et al., 2020). Thus, the parcellation of neural activity space into covariation patterns that subserve distinct functions may reflect a general principle of neural systems.

## Conclusion

Using simultaneously recorded motor cortical and muscle activity in a new behavioral paradigm, we found that the specificity of neurons for antagonist cocontraction or alternation tasks exists along a continuum rather than in discrete groups, that cocontraction is not associated with a suppression of intracortical inhibition, and that neuronal populations do not linearly encode muscle activity consistently across tasks. These results contradict existing views of motor cortical operation that are grounded in muscle- or task-based cellular organization. Instead, our results indicate that rather than relying on coactive flexion and extension programs or the contributions of specialized cell classes, tasks comprising different muscle activation are driven by different motor cortical activity covariation, which we observed for corticospinal neurons specifically. This aligns with the emerging view that specific motor functions may be mediated by distinct covariance patterns in motor system activity.

### Limitations of the study

Our study has several limitations. In particular, our results do not directly address what it is that causes the differences in activity covariation we observe between tasks. There are numerous potential causes, including sensory inputs, and further study would be necessary to define relevant inputs and aspects of local activity dynamics. We are also unable to directly link the observed changes in activity covariation to alternation and cocontraction of forelimb muscles per se, to the exclusion of other aspects of motor output involved in task performance. The muscle activation patterns during the two tasks do not differ solely in the degree of alternation and cocontraction, but also in other aspects of their temporal pattern. Additionally, due to the relative difficulty of data collection, we analyzed electrophysiological data from four male mice and calcium indicator imaging data from two male mice; data from additional subjects would further bolster our findings.

## Supporting information

Movie S1 alternation task (0.5 speed)

Movie S2 cocontraction task (0.5 speed)

## ACKNOWLEDGMENTS

We are grateful to M. Marshall for illustrations; C. Schoonover, A. Fink and J. Climer for technical advice; M. Mendelsohn and N. Zabello for animal care; E. Famojure, B. Han, K. MacArthur, M. Marshall, I. Shieren, G. Martins, H. Rodrigues, and M. Correia for technical assistance; and T.M. Jessell for invaluable early scientific guidance when the project was devised. C.L.W. was supported by a T32 and F31 NIH-NINDS Training Grant. A.M. is supported by a Searle Scholar Award, a Sloan Research Fellowship, a Whitehall Research Grant Award, the Chicago Biomedical Consortium with support from the Searle Funds at the Chicago Community Trust, and NIH grant DP2 NS120847.

## AUTHOR CONTRIBUTIONS

C.L.W. and A.M. devised the project and designed experiments. C.L.W., S.F., and A.M. performed the experiments. C.L.W. and A.M. analyzed data with assistance from S.S. All authors helped interpret data. C.L.W. and A.M. wrote the manuscript, with input from R.M.C.

---

**Movie S1. Mouse performing the alternation task, related to Figure 1**

Video depicts a head-fixed mouse performing the alternation task. The mouse reaches forward with the right forearm to grasp the rungs of a small wheel and pulls it. This wheel pulling involves alternating contractions in triceps and biceps. After a target distance is achieved, reward is dispensed in the form of water droplets; the number of droplets reflects the speed at which the mouse pulled the wheel relative to the previous 10 instances to encourage faster and more stereotyped movement. An LED seen at bottom left flashes each time a droplet is administered. Video was originally recorded at 60 fps but is shown here at 0.5x actual speed.

**Mouse S2. Mouse performing the cocontraction task, related to Figure 1**

Video depicts a head-fixed mouse performing the cocontraction task. The mouse rests its paw on a handle, though is not required to do so, and generates transient triceps-biceps cocontraction for water reward. Muscle activity is analyzed online: if normalized triceps and biceps activation were suprathreshold, concurrent, and within a certain range of each other, reward is dispensed. The number of water droplets dispensed is dependent on the correlation of smoothed triceps and biceps signals relative to the previous 10 instances to encourage increased correlation. An LED seen at bottom left flashes each time a droplet is administered. Video was originally recorded at 60 fps but is shown here at 0.5x actual speed.

## METHODS

### Key resource table information

#### Chemicals, Peptides, and Recombinant Proteins

- Muscimol hydrobromide, Sigma, Cat. #: G019-5MG
- retro-hSyn1-GCaMP6f, HHMI viral core (Tervo et al., 2016)

#### Experimental Models: Organisms/Strains

- Mouse: C57BL/6J, The Jackson Laboratory, Jax stock #: 000664; RRID: IMSR_JAX:000664
- VGAT-ChR2-EYFP, B6.Cg-Tg(Slc32a1-COP4*H134R/EYFP) 8Gfng/J, The Jackson Laboratory, JAX stock #: 014548; RRID: IMSR_JAX:014548
- GAD2-IRES-Cre, (C57BL/6 x 129S4/SvJae)F1, The Jackson Laboratory, JAX stock #: 010802

#### Software and Algorithms

- MATLAB v.9.4.0 (2018a), MathWorks, https://www.mathworks.com/products/MATLAB/
- KlustaKwik, Rossant et al., 2016, http://klusta.readthedocs.io/en/latest/

#### Other

- Cerebus Neural Signal Processing System, Blackrock Microsystems, 128 channels
- Prairie Technologies two-photon microscope and data acquisition system
- 4-channel EMG amplifier, University of Cologne Electronics Lab, Model #: MA 102S
- PCIe-6323 DAQ, National Instruments, Cat. #: 781045-01
- Laser displacement sensor, Micro-Epsilon, Cat. # optoNCDT 1302 ILD 1302-200
- Linear actuator, Actuonix, Cat. # L12-50-50-12-S
- Silicon probe, NeuroNexus, Cat. #: A1x32-Poly3-5mm-25s-177
- Stainless steel wire for EMG electrodes, A-M Systems, Cat #: 793200

### EXPERIMENTAL MODEL AND SUBJECT DETAILS

All experiments and procedures were performed according to NIH guidelines and approved by the Institutional Animal Care and Use Committee of Columbia University.

#### Experimental Animals

A total of 54 adult male mice were used, including those in early experimental stages to establish methodology. Strain details and number of animals in each group are as follows: 6 VGAT-ChR2-EYFP line 8 mice (B6.Cg-Tg(Slc32a1-COP4*H134R/EYFP) 8Gfng/J; Jackson Laboratories stock #014548); 11 GAD2-IRES-Cre mice (Taniguchi et al., 2011; (C57BL/6 x 129S4/SvJae)F1; Jackson Laboratories stock #010802); and 37 C57BL/6J mice (Jackson Laboratories stock #000664).

All mice used in experiments were individually or pair housed under a 12-hour light/dark cycle. At the time of the measurement reported, animals were 10–20 weeks old. Animals weighed approximately 23-28 g. All animals were being used in scientific experiments for the first time. This includes no previous exposures to pharmacological substances or altered diets.

### METHOD DETAILS

#### Alternation and Cocontraction Task

Male mice were trained through a behavioral shaping procedure to perform, within a single behavioral session, three epochs of behavior: first an alternation task, then a cocontraction task, and finally a second epoch of the alternation task. During the alternation task, mice used their right forelimb to repeatedly pull the rungs of a wheel towards them, eliciting alternating triceps and biceps muscle contractions (Figure 1D-G-D). After 150 µl of water had been earned, a platform automatically extended to block access to the wheel and the cocontraction epoch began. During the cocontraction task, mice co-activated their triceps and biceps muscles after a period of muscle quiescence (Figure 1H-K-H). After another 210 µl of water reward had been earned, the platform was automatically retracted, and a second epoch of the alternation task takes place until a final 150 µl of reward had been dispensed.

##### Apparatus

Head-fixed mice were placed within an enclosure consisting of a flat platform (10 cm by 6 cm) and a curved hutch (4.5 cm outer diameter, 4.8 cm length) lined with soft foam (3 cm inner diameter) that covered the posterior half of their body. The front-right quadrant of the platform was removed, allowing a wheel (60 mm diameter, 11.5 mm width) bearing 28 rungs (1.4 mm diameter) to be placed within reach of the right forelimb for use during the alternation task. A rounded barrier below the mouse’s chest prevented mice from manipulating the wheel with their left forelimb. Platform, hutch, and wheel were custom designed and 3D printed using metallic plastic (Shapeways).

The behavioral assay was controlled using the MATLAB Data Acquisition Toolbox and the NI PCIe-6323 DAQ. The wheel was affixed to an 8 in. stainless steel shaft whose ends were mounted on bearings (8600N1, McMaster-Carr). An absolute rotary encoder was mounted around the shaft and measured its angular position (A2K-A-125-H-M, U.S. Digital). A ratchet mechanism ensured that the wheel could only rotate towards the mouse. A one-dimensional laser displacement sensor was positioned in front of the mouse and aimed at the midpoint of the right forelimb to detect its position along the anterior-posterior axis (optoNCDT 1302, Micro-Epsilon). When the behavior epoch transitioned from the alternation task to the cocontraction task, a custom platform (5 by 7 cm) bearing a handle identical to the wheel’s rungs, was automatically positioned over the wheel by a linear actuator (L12-50-50-12-S, Actuonix). Water rewards (3 µl/droplet) were dispensed with a solenoid valve (161T012, NResearch) attached to a lick-port (01-290-12, Fisher). Mice were exposed to static visual cues, displayed on a computer monitor about 1 foot from the headplate holder, that corresponded to the task they were meant to perform: horizontal black bars on a white background during alternation and vertical bars during cocontraction. However, these visual cues were dispensed with during calcium imaging due to the potentially deleterious effect of visible light on the functional signal.

##### Electromyographic Electrode and Headplate Implantation

Electromyographic (EMG) electrodes were fabricated for forelimb muscle recording using a modification of established procedures (Akay et al., 2006; Pearson et al., 2005). One pair of electrodes was used per muscle recorded and electrode sets were made up of multiple pairs of electrodes. Each pair was composed of two 0.001-inch braided steel wires (793200, A-M Systems) knotted together. For one wire of each pair, insulation was stripped from 1 mm to 1.5 mm away from the knot. On the other, insulation was stripped from 2 to 2.5 mm away from the knot. The ends of the wires above the knot were soldered to a miniature connector (11P3828, Newark). The length of wire left between the knot and the connector depended on the proximal-distal location of the muscle targeted for recording: 3.5 cm for pectoralis, spinodeltoidus, biceps and triceps (lateral head), and 4.5 mm for extensor digitorum communis and palmaris longus. For the functional imaging experiments, only the last four of these muscles were recorded. Insulation was removed from the remaining ends of the pairs of wire, which were then twisted together and crimped within a 27-gauge needle that allowed insertion into the muscle.

Under anesthesia induced by isoflurane (1-3%; Henry Schein), the neck and right forelimb of the mouse was shaved, and incisions were made above the muscles to be implanted. Electrode pairs were guided from the incision at the scalp, under the skin, to the incisions at the forelimb. Electrodes were inserted into muscle using the needle, after which a knot was made in the distal portion of the electrodes to secure them, and the excess wire and needle were cut away. The incisions in the forelimbs were closed with sutures.

Within the same surgery, titanium headplates (25 x 9 x 0.8 mm) were affixed to each mouse’s skull using dental cement (Metabond, Parkell). The center of the headplates was open, allowing subsequent access to the skull covered with dental cement. Upon headplate implantation, the location of bregma relative to a mark on the headplate was recorded to facilitate the positioning of craniotomies during subsequent surgeries. After headplate implantation, the EMG electrode connector was attached to the posterior edge of the headplate using dental cement and the incision at the back of the scalp was closed with sutures. Analgesia in the form of subcutaneous injection of carprofen (5 mg/kg) and bupivacaine (2 mg/kg) were delivered during surgery. Carprofen was again administered at 24- and 48-hours post-surgery. After recovery from the implantation surgery, mice were placed on a water schedule in which they received 1 mL of water per day.

##### EMG Recordings

Electromyographic recordings were amplified and bandpass filtered (250-20,000 Hz) with a differential amplifier (MA102 with MA103S preamplifiers, University of Cologne electronics lab). Data were digitized and acquired at 30 kHz using the Cerebus neural signal processing system and Central Software Suite (Blackrock Microsystems).

Before behavioral training, EMG recordings were made while mice ran on a custom-built, motor-driven rodent treadmill (Model 802, University of Cologne electronics lab) at a rate of 20 cm/s. We verified that EMG measurements reflected muscle activity instead of motion artifact through the presence of spike-like transients with alternating activation and quiescence during locomotion. However, we note that we were not able to exclude the possibility that EMG recordings were influenced by the activity of nearby muscles. Using the recordings made during locomotion, the 0.01th and 99.9th percentile value was calculated for each EMG channel and used to normalize subsequent EMG recordings during the behavioral paradigm as percentages of maximum muscle contraction observed during locomotion.

##### Training

Four days after the beginning of the water schedule, mice were acclimated to being handled by the experimenter following established protocols (Guo et al., 2014b). After two daily sessions of acclimation to handling, mice were acclimated to being head-fixed on the behavioral apparatus for 15 minutes on the first day and 30 minutes on the second day, during which time they were provided with water rewards (3 µl per reward) at regular intervals. During acclimation, the wheel was locked in place to prevent its movement.

After acclimation, mice underwent twice daily 30–45-minute training sessions of the behavioral paradigm. The 3-week behavioral shaping procedure involved a first week during which mice were trained in the alternation task exclusively, followed by a week of training in the cocontraction task exclusively, and finally a week during which mice would be trained to perform both tasks within one behavioral session. During the first training session of the alternation-only week, the wheel was unlocked, and rewards dispensation was triggered by the experimenter’s keypress whenever the mouse placed its paw on the wheel and/or performed any slight rotation of the wheel towards itself. Over the course of this first session, mice learned to associate wheel rotation with reward and began iteratively pulling the rungs of the wheel. Over the following ∼13 sessions, mice were gradually trained to pull the wheel with increasing rapidity. The distance of wheel rotation was integrated in behavioral software until a certain threshold was achieved, the time to reach the threshold distance was determined, and the integrated distance was reset to 0. Mice automatically received a water reward for each of the first 10 instances in which the distance threshold was met. For subsequent instances, the time to reach threshold was compared to those from the previous 10 instances. If that time was below the 10^th^ percentile value, 4 rewards were dispensed. If the time was below the 40^th^ percentile value, 2 rewards were dispensed, and if the time was below the 75^th^ percentile value of those times, one reward droplet was dispensed. If the time was above the 75^th^ percentile value, no rewards were dispensed. Every minute, the distance threshold was adaptively updated to maintain reward rate at 0.5 mL of water dispensed per training session; if the reward rate was too low, the distance threshold was lowered, and if the rate was too high, the threshold was raised.

During the following ∼14 sessions, mice were gradually trained to maintain triceps and biceps quiescence (<10% maximum contraction) for a minimum of 500 ms before concurrently contracting the triceps and biceps muscles at increasing amplitudes, durations, and levels of triceps-biceps activity correlation. For this training, a platform bearing a handle identical to the wheel’s rungs was moved into place above the wheel to prevent access to it. During this task, triceps and biceps activity was processed using the behavioral software and monitored for cocontraction events. During the first cocontraction training sessions, triceps and biceps were required to contract above an amplitude threshold of 15% of maximum contraction for a duration of at least 32 ms. The amplitude threshold and duration threshold were increased manually at the beginning of each session until a final threshold of 40% and at least 160 ms, respectively, were achieved. Mean normalized triceps and biceps activity was also required to be within a certain range of each other, starting with the requirement that neither be more than double the other. This range was reduced manually over training down to a 1.25-fold difference. Within a session, when the conditions for a cocontraction event were met, the correlation of the envelope of triceps and biceps activity was calculated for 300 ms surrounding the time of amplitude threshold crossing. Mice automatically received water reward for the first 10 cocontraction events. After this, the triceps-biceps correlation was compared to the previous 10 instances. If the correlation was greater than the 90^th^ percentile, 4 rewards were dispensed, if the correlation was greater than the 60^th^ percentile, 2 rewards were dispensed, and if the correlation was greater than the 25^th^ percentile, 1 reward droplet was dispensed. If the correlation was below the 25^th^ percentile value, no rewards were dispensed.

To generate the behavioral learning curves and to perform the muscimol inactivation experiments (Figure 1L-Q-O), one cohort of 5 mice was used. Mice were trained in twice-daily sessions of only the alternation task for one week (Figure 1L), then underwent muscimol perturbation during alternation (described below) over the course of 4 days. Mice were given a week with no training before starting twice-daily sessions of only the cocontraction task over the course of the subsequent week (Figure 1M), after which they underwent the muscimol perturbation during cocontraction. The tasks’ performance sensitivity to muscimol were assessed individually in order to isolate their respective cortical dependence in the absence of the difficulty inherent in learning to perform two different tasks within the same session.

#### Muscimol Injection

Muscimol perturbations took place immediately following a week of twice-daily training sessions in either the alternation task or the cocontraction task. A day prior to the beginning of injections, the dental cement covering the skull was drilled away and a 1 mm diameter craniotomy was made above the left CFA. After making the craniotomy and following each round of injections, craniotomies were sealed with Kwik-Cast (WPI). Injections were performed 90 minutes before the 2^nd^ training session of the day. A Nanoject II (Drummond) equipped with pulled glass capillaries was used to inject 1 ng/nl muscimol hydrobromide (G019-5MG, Sigma) in saline (DPBS with CaCl_2_ and MgCl_2_, GIBCO) or saline alone. Injections were made in the center of the CFA, as previously delineated (1.5 mm left and 0.25 mm rostral of bregma; Tennant et al., 2011). Two boluses of 36.8 nl were extruded: one at 700 µm and one at 400 µm below the pial surface. Extrusion was verified to be successful immediately before and after the insertion of the capillary into neural tissue. A wash-out behavioral session was performed the following morning. Contralateral injections were performed over 2 days (Figure 1N-Q-O) and bilateral injections were performed over the following 2 days (Figure S1I, M, L). The cohort of 5 mice was split into groups of 2 and 3 mice such that one group received muscimol injection while the other received saline one day, and vice versa on the following day.

#### Electrical Recording in Motor Cortex

The day before recordings began in trained animals, dental cement above the skull was removed and a square craniotomy (1.5 x 1.5 mm) was made over the left CFA with a surgical drill and scalpel. The exposed tissue was covered with Kwik-Cast. To implant bone screws, two ∼0.7 mm diameter craniotomies were made over the right parietal and occipital cortices. A #000 screw (B000FN0J58, Amazon) soldered to a male connector pin (520200, A-M Systems) was positioned in each craniotomy and rotated until in contact with the brain surface. Dental cement was then applied to the exposed skull and screw, leaving the male connector pins exposed. During recording, female connector pins (520100, A-M Systems) soldered to both ends of 2-stranded ribbon wire (10647, SparkFun) served to connect the bone screws to the male connectors on a 32-electrode silicon probe (A1x32-Poly3-5mm-25s-177, NeuroNexus).

Before recording, the trained animal was head-fixed on the behavioral apparatus and the Kwik-Cast removed from the craniotomy, exposing the surface of the brain. Probes were slowly lowered to a depth of 650-800 µm from the pial surface at an approximate rate of 2 µm/s and allowed to settle in the tissue for 20 minutes before behavior and neural recording were initiated. Neural activity was acutely recorded using a digital headstage (Cereplex M64, Blackrock Microsystems). Data was acquired at 30 kHz using the Cerebus 128-channel neural acquisition system and Central software (Blackrock Microsystems).

#### Optical tagging

To determine the relationship between waveform width and inhibitory interneuron identity, we performed optical tagging in VGAT-ChR2-EYFP mice. Untrained mice received a 2-2.5 mm diameter craniotomy over the left CFA and bone screws over the right parietal and occipital cortices, as described above. Before recording, the animal was head-fixed and the Kwik-Cast removed, exposing the surface of the brain.

The 32-electrode silicon probe was slowly inserted at a 45° angle to vertical depths of 800, 1000, or 1200 µm from the brain surface and allowed to relax for 20 minutes. A 2.5 mm ceramic ferrule (M81L01, Thorlabs) connected to a 200 µm core, 0.39 NA optical patch cable was positioned above the craniotomy so that a 2 mm diameter spot of light projected onto the surface of the brain from a 473 nm laser (CL473-075-O, CrystaLaser). Pulses of light at an intensity of 10 mW/mm2 were applied to the brain surface at a frequency of 20 Hz and a duty cycle of 50%. Light intensity and duty cycle were set according to experiments calibrating the relation between light power and cessation of pyramidal neuron activity due to the activation of inhibitory interneurons (Guo et al., 2014b). Neural activity was recorded as described above.

#### Spike Sorting

Putative neural spikes were detected and sorted using KlustaKwik (Rossant et al., 2016). Clusters of waveforms were categorized as corresponding to well-isolated units if their spike autocorrelograms showed an absolute refractory period of a minimum of 1 ms and a firing rate far less than the unit’s mean firing rate for a minimum of 2 ms before and after a spike occurred. Any clusters that did not meet these criteria were discarded. If more than one cluster contained spikes for the same unit, as demonstrated by the presence of similar waveforms across clusters and the absence of refractory period violations in their cross-correlograms, the clusters were merged. If a single cluster contained activity from more than one unit, as demonstrated by the presence of dissimilar waveforms or refractory period violations, the cluster was manually split.

#### Parameters used and descriptions from KlustaKwik documentation

(Note: the int() function returns the nearest integer, and text written after “#” is commented out) Bit depth of raw data: nbits = 16

Multiplier from actual voltage to stored data: voltage_gain = 10.

Raw data sampling rate, in Hz: sample_rate = 30000

Number of channels in the recording: nchannels = 32

Bandpass filter low corner frequency: filter_low = 500.

Bandpass filter high corner frequency: filter_high = 0.95 * .5 * sample_rate

Order of Butterworth filter: filter_butter_order = 3

Raw data chunk size: chunk_size = int(1. * sample_rate) # 1 second

Overlap of chunks: chunk_overlap = int(.015 * sample_rate) # 15 ms

Number of uniformly distributed chunks used to estimate its standard deviation: nexcerpts = 50 Length of chunks (seconds): excerpt_size = int(1. * sample_rate)

threshold_strong_std_factor = 4.5

threshold_weak_std_factor = 2.

The number of samples to extract before and after the center of the spike for waveforms: extract_s_before = 16 extract_s_after = 25

Number of features (PCs) per channel: nfeatures_per_channel = 3

The number of spikes used to determine the PCs: pca_nwaveforms_max = 10000 Number of samples to use in floodfill algorithm for spike detection:

connected_component_join_size = 1

connected_component_join_size = int(.00005*sample_rate)

Waveform alignment: weight_power = 2

Whether to make the features array contiguous: features_contiguous = True

#### Two-photon imaging

##### Spinal injections

For the calcium imaging experiments, GAD2-IRES-Cre (Taniguchi et al., 2011) mice underwent cervical spinal injection surgery (Azim et al., 2014; Fink et al., 2014) the day prior to EMG electrode and headplate implantation surgery. Under anesthesia induced by isoflurane, the neck and upper back of the mouse was shaved, and an incision was made over the cervical spinal cord vertebrae. The muscle overlying the area was gently pushed away and the T2 spinous process was secured and slightly lifted to better expose the cervical vertebrae. Dura was removed from between the vertebrae. A Nanoject II/III equipped with a pulled glass capillary was used to inject multiple boluses of 13.8 nl of AAV2-retro-hSyn1-GCaMp6f (HHMI Viral Core, (Tervo et al., 2016) and 13.8 nl of EF1α-FLEX-TVAmcherry (Watabe-Uchida et al., 2012) and AAV2/1-CAG-FLEX-H2B-GFP-2A-CVS-N2cG (Reardon et al., 2016) unilaterally at two sites in each brachial segment from C4 to T1. These last two “helper” AAVs permit expression of an avian receptor A (TVA) and rabies glycoprotein (G) under the control of Cre. In this way, the infection and monosynaptic transport of the glycoprotein deleted rabies is restricted to GAD2+ spinal interneurons. After functional imaging was complete, a 2^nd^ spinal injection surgery took place in which the modified glycoprotein-deleted rabies virus RABV-N2C^ΔG^-EnvA-tdTomato was injected in a manner identical to the initial spinal surgeries. After 10 days, we examined rabies expression in CSNs that contact GAD2+ cells.

##### Cranial window implantation

Behaviorally trained mice underwent cranial window implantation surgery a minimum of three days before recording of neural activity began. Cranial windows were constructed from two laser-cut, truncated half-circle shaped glass window (2.5 x 1.5 mm at widest points, 200 µm thick, Tower Optical Corp.) adhered to a larger 4 mm diameter (No. 1, Warner Instruments) glass coverslip using UV-curing adhesive (Norland Optical Adhesive 61). Under anesthesia induced by isoflurane and after administration of the anti-inflammatory drug dexamethasone (2 mg/kg), a surgical drill was used to remove the dental cement covering the skull. The surgical drill and a scalpel were used to create a rectangular craniotomy (∼1.5 by 3 mm) centered at the CFA using the fiducial on the headplate for bregma coordinates. Dura was gently removed using fine, angled forceps and a micro-surgical knife. The cranial window was lowered into place using a custom vacuum system (suction through an unpulled glass capillary tube) until the entirety of the smaller glass surface was in contact with brain tissue. The larger glass surface rested against the top of the skull and was affixed using dental cement around its perimeter. The surface of the cranial window was protected with a layer of Kwik-Cast that was later removed before imaging.

##### Imaging

Two-photon fluorescence imaging was carried out using a 25X 1.0NA objective (Olympus) mounted on Ultima microscope (Prairie Technologies) and using a Ti:sapphire laser (Chameleon Ultra II, Coherent) tuned to 940 nm (GCaMP6f) or 1050 nm (tdTomato). Images were acquired using the PrairieView 5.4 software. GCaMP6f image time series were acquired at a rate of 60 Hz, and every four images were averaged for a final sampling rate of 15 Hz. Images covered ∼296.5 x 296.5 µm with 256 x 256 square pixels. Muscle activity and behavioral parameters were collected (General Purpose Input Output (GPIO) Box, Prairie Technologies) at a sampling rate of 10 kHz also via the PrairieView 5.4 software, allowing for alignment of imaging with behavioral data. GCaMP6f image time series was collected for each animal during 20-40 minutes of behavioral performance for up to 7 daily behavioral sessions. Ten days after rabies injection, Z image stacks were collected over 40-80 µm depths with 1 µm separating each image both for GCaMP6f and tdTomato in each plane. The Z ranges spanned the expected plane of previous GcaMP6f image time series collection. The red emission collection channel registered no detectable signal at 940 nm excitation, indicating that red emission was generated solely by tdTomato and not GcaMP6f.

### QUANTIFICATION AND STATISTICAL ANALYSIS

All analysis was completed in MATLAB v.9.4-9.9 (MathWorks).

#### EMG Processing and Analysis

EMG measurements were low-pass filtered and downsampled to 1 kHz (MATLAB function ‘decimate’ which uses a low-pass Chebyshev Type I infinite impulse response (IIR) filter, order = 8) then high-pass filtered at 40 Hz (Butterworth IIR, order = 12), full-wave rectified, and convolved with a symmetric Gaussian that had a 10 ms standard deviation.

##### EMG during Alternation and Cocontraction Training and Muscimol Inactivation

One cohort of 5 mice was used to quantify learning curves and for the muscimol inactivation experiments. To normalize EMG activity during these behavioral sessions, the 0.01^th^ and 99.99^th^ percentile values were taken as minimum and maximum values, respectively, from the three baseline sessions preceding the onset of the perturbation experiments. These values, which were highly similar to those found during treadmill locomotion, were averaged across the three sessions and used to normalize each muscle’s activity. Due to variability in behavioral session length, only the first 20 minutes of each session were analyzed.

Task performance was measured using an automated process. During the alternation task behavioral sessions, task performance was measured as mean wheel velocity. This measure was calculated from the absolute encoder signal mounted around the axis of the wheel. During the cocontraction task behavioral sessions, task performance was measured as the number of cocontraction events. To assess this parameter, the cocontraction index (CCI) between triceps and biceps was calculated for each timepoint by dividing the amplitude of the less active muscle by that of the more active muscle, then multiplying the product by their sum (Knarr et al., 2012; Rudolph et al., 2000). Peaks in the CCI above a threshold of 0.2 were used to identify segments of interest, and these segments were aligned on the rising edge of triceps activity (the time at which 50% of peak amplitude attained before peak). To eliminate segments in which cocontraction did not begin from a period of muscle quiescence, any segments in which the CCI went above a threshold of 0.1 in the period from 150 to 100 ms preceded the alignment point were eliminated. Remaining events were retained and counted as cocontraction events. Though during cocontraction training the amplitude threshold above which a cocontraction event is rewarded is adaptive, analysis of learning and response to muscimol perturbation was performed using the fixed threshold of 0.2 CCI to enable comparison across time and conditions.

##### EMG during Alternation and Cocontraction Behavior and Neural Recording

Analysis of EMG recorded during neural recording required an automated identification of alternation and cocontraction trials that were matched for triceps and biceps amplitude within-session. During alternation, candidate trials were identified by the presence of antiphasic triceps and biceps activity. During cocontraction, candidate trials including cocontraction events were identified by the presence of highly correlated, suprathreshold triceps and biceps activity preceded by muscle quiescence. For both tasks, trials were aligned on the rising edge of triceps activity. For each recording session, 30 trials in each behavioral epoch were sub-selected algorithmically based on triceps-biceps correlation and differences in their amplitudes in order to maximize within-epoch similarity and to match muscle activity levels across tasks. The timestamps of these trials were then used to identify concurrent limb position and activity of other recorded muscles. The activity of the other recorded muscles during alternation trials (Figure S1B, C) and cocontraction trials (Figure S1D, E) was also compiled.

Each muscle’s activity was normalized by its minimum (0.01^th^ percentile) and maximum (99.99^th^ percentile) value taken from the entire behavioral session, which were similar to those recorded during treadmill locomotion. First, potential alternation and cocontraction trials within a session were identified from each epoch based on triceps and biceps activity. During the alternation epochs, time segments containing triceps activity peaks above a set threshold were identified. If within a 10 ms window around that peak, biceps activity was below a set threshold (indicating that there was low or no biceps activity during triceps contraction), the time segment was retained as a potential alternation trial. During the cocontraction epoch, time segments containing triceps activity peaks above a set threshold were identified. If biceps activity peaked above that same threshold in a 50 ms window around the triceps peak, the time segment was retained as a potential cocontraction trial. After potential trials were identified, 30 trials from each epoch in each session were subselected based on muscle amplitude and trial quality. First, trials were aligned at triceps activity onset (at 50% of the triceps peak amplitude) then assessed based on 4 parameters. For alternation, these parameters were the triceps peak amplitude, the biceps peak amplitude, the difference between these two amplitudes, and the triceps and biceps correlation (which during alternation was expected to be low). For cocontraction, these parameters were the triceps peak amplitude, the difference between the triceps peak amplitude and the biceps amplitude at that time point, the muscle activity during the quiescent period preceding cocontraction, and the Euclidean distance between the triceps and biceps activities. Hard thresholds were set for each parameter and any trials that did not meet these thresholds were eliminated. Target values were also set for each parameter, and the remaining trials received a score based on the distance between their metrics and the target such that a low score indicated a high-quality trial. A combination of weights was then iteratively applied to each metric’s score and, at each iteration, the 30 trials per epoch that had the lowest score were chosen. The selected trials were then evaluated for respective triceps and biceps variability within-epoch, root mean squared error of respective triceps and biceps activity across the first and second alternation epochs, and the difference in peak muscle amplitudes within-epoch and across all three epochs. The set of trials that had the lowest evaluation score was then selected for that session.

Periods of muscle quiescence were identified by examining the activity of all 6 recorded muscles. Starting at a threshold of 0.015 normalized amplitude (% of maximum contraction), the number of 500 ms time segments during which each muscle’s activity was below this threshold. If fewer than 90 segments were found, the threshold was incrementally increased by 0.001 normalized amplitude until this condition was met. Then, every 3rd segment was selected, to ensure better coverage of the session in time, and trimmed to 300 ms, resulting in 30 trials of muscle quiescence.

Finally, the quality of each behavioral session was evaluated. A session was retained for analysis if: the correlation of triceps activity in alternation 1 and 2 and the correlation of biceps activity in alternation 1 and 2 each had a coefficient greater than 0.5; the correlation of triceps and biceps activity in alternation 1, and separately in alternation 2 had a coefficient less than −0.5; and the Euclidean distance between triceps and biceps activity during cocontraction was less than 0.9. Six to eight recording sessions per mouse included behavior of sufficient quality to be retained for analysis.

To assess the generalization of our findings to wrist alternation and cocontraction, a light-weight trial selection was carried out based on extensor digitorum communis (EDC) and palmaris longus (PL) activity from the behavioral sessions selected for analysis by the procedure described above. Peaks of EDC activity, normalized as described above, of at least 0.2 were identified. For cocontraction trials, any trial in which normalized PL in a 20 ms window around the EDC peak was 0.2 or higher were retained. The trials per session were then aligned on EDC contraction onset at 50% peak height. The Pearson’s correlation coefficient for EDC and PL activity was determined for the 100 ms window around EDC onset, and no more than the 50 most highly correlated trials were retained. The Euclidean distance was then taken over the same window, and the 30 trails with the lowest distance were retained for further analysis. Similarly, for alternation trials, only trials in which PL activity was below 0.2 in the 20 ms around the EDC peak were retained, followed by alignment of trials on 50% EDC onset. Subsequently, no more than 50 trials with the lowest correlation between EDC and PL in the 100 ms around EDC onset were retained, followed by the 30 trials with the highest Euclidean distance. Using this procedure, in all 4 mice EDC-PL alternation could be found, but only 3 of 4 mice reliably performed EDC-PL cocontraction in each behavioral session. A36 only exhibited cocontraction in 2 of 6 sessions and was therefore not included in analysis of either wrist cocontraction or alternation.

#### Firing Rate Calculation

Firing rates were calculated at 1 kHz for each well isolated unit by defining for each spike a time series approximating a Gaussian function with its mean at the time of the spike and a standard deviation of 10 ms, and which was normalized to temporally integrate to 1. These time series were then summed, producing a smoothed firing rate time series. All plotting and analysis of neural data, save the subtype assignment in the optical tagging experiments, used these smoothed firing rate time series. Trial averages for behavioral epochs were assembled using these time series.

Neurons that had a firing rate below 1 Hz in all behavioral epochs and neurons that had an alternation 2 mean firing rate greater than its mean alternation 1 firing rate multiplied by 1.5 or less than its mean alternation 1 firing rate divided by 1.5 were excluded from analysis due to inconsistent task-relatedness. Modest adjustments of these parameters did not change overall results. Additionally, forgoing the step of removing neurons due to inconsistent task-relatedness had no significant change to the results. To control for probe drift, any neurons whose initial spike waveform (as averaged over the first 300 spikes) varied dramatically from later in the session was excluded. This was carried out by calculating the Euclidean distance (MATLAB function ‘pdist’) of the initial waveform to all subsequent waveforms. If the slope of the resulting absolute distance values was greater than 0.2, the neuron was not included in analysis. Pearson’s correlations between firing rates and the summed time series of each muscle’s activity, both further smoothed with a moving window of 1 s, were computed using the MATLAB function ‘corr’ (Figure 2I).

#### Waveform-Based Subtype Assignment

Distributions of waveform widths were separately determined for the dataset of neural activity recorded from trained mice during behavior and for the dataset of neural activity recorded during optical tagging. Once the optimal classification threshold had been determined for the distribution of widths from the optical tagging data, this threshold was applied to the distribution of widths from the dataset recorded during behavior to identify putative inhibitory interneurons and putative pyramidal cells.

For each isolated unit, the electrode on which the spike-related voltage transient had the largest amplitude was identified. The mean spike waveform was computed as the mean voltage time series on the identified electrode from 1 ms before the spike to 1.5 ms afterward over a maximum of the first 300 recorded spikes. The trough-to-peak spike width was measured as the time between the mean spike waveform’s minimum and its subsequent maximum.

For the optical tagging dataset, neurons were identified as light-responsive if the latency of its first spike in the 10 ms after onset of light stimulus was shorter than would be expected by chance (p<0.05), as compared to stimulus-free timepoints, over the course of 148 trials (Stimulus-associated spike latency test [SALT], (Kvitsiani et al., 2013). Given the distribution of the tagged cells relative to untagged cells, the optimal waveform width threshold of 0.42 ms was determined by finding the point at which the ratio of the fraction of tagged cells among the narrow-waveform group to the fraction of tagged cells among the wide-waveform group was maximal. At this threshold, 63% (17/27) of the narrow-waveform neurons and 11.5% (9/78) of the wide-waveform neurons were optically tagged and thus identified as GABAergic. This optimal classification threshold was then applied to the distribution of waveform widths recorded during behavior and allowed assignment of neurons to either narrow-or wide-waveform groups

#### Linear Regression of muscle activity to neural activity

Ridge regression was used to fit a model in which the activity of each muscle is determined by neural firing rates separately during alternation and cocontraction behavior. The trial-averaged activity of each muscle was mean-centered and fit by a linear combination of the trial-averaged firing rates. For matrices *M* of muscle activity and *N* of firing rates, in which each row corresponds to a different time point and each column to a different muscle or neuron, the model is defined as *M = NW* where *W* is a matrix of weights and is in the absence of any temporal offset. For each mouse, neurons were fit to a model that was compared to a mean of each muscle’s activity across all sessions.

For each epoch (excluding muscle quiescence), sets of 30 trials were randomly partitioned into three equal subsets (train, validate, and test) and activity was averaged within each subset. The ridge parameter was selected using a cross-validation procedure. A range of parameter values was used to train the model on the train subset then subsequently tested on the validate subset. The parameter value that yielded the highest R^2^ value (coefficient of determination) was chosen. The train and validate subsets were then averaged, the chosen ridge parameter used when training the weights on the averaged activity, and these weights were tested on the test subset within that epoch and on the test subsets in other epochs. For each mouse, this procedure was repeated for 500 permutations of trials among the subsets and the resulting R^2^ values were averaged. The set of weight vectors that produced the model whose R^2^ value was closest to the mean R^2^ across all permutations was retained to provide example model fits. The joint regression analysis was performed identically save that the alternation 1 and cocontraction activity were concatenated in time.

The linear models trained on each epoch type were used to extract muscle-related neural signal. The 500 sets of weight vectors that produced the model trained on each epoch were averaged and orthonormalized using the Gram-Schmidt process, and the projection of the neural data *N* onto these weights was computed. This projected data was multiplied by the transpose of the orthogonalized weight matrix yielding a matrix of only the neural signals that contributed to the modeling of the muscle signals.

#### Analysis of Weighted Sums Defined by Principal Components

In Figure 5A-E, principal components (PCs) were calculated separately for each behavioral epoch using matrices of trial-averaged neuronal firing rates, *D*, each having rows corresponding to timepoints and columns corresponding to neurons, using the “pca” function in MATLAB. Six PCs were determined for combined alternation and cocontraction data by concatenating the behavioral epochs in time before mean-centering their activity. Projections onto individual principal components were determined by multiplying *D* by the column vector composed of the weight on each cardinal neural dimension for a given component. To compute the normalized variance of other epochs in the alternation 1 epoch, the dimensionality of trial averaged data was reduced to 6 by projecting it onto the basis sets of their top 6 PCs. Normalized variance for each segment was then computed as the trace of the covariance matrix for each of these projected segments, normalized by the trace of the covariance matrix for the full projection matrix of activity during alternation 1. This normalization by the total variance during alternation 1 allows a comparison of magnitudes across tasks.

The alignment index, measuring alignment between neural activity during the first epoch of alternation and the other behavioral epochs, was computed for sets of trial-averaged firing rates. For the alternation 1 dataset, *D_1_*, and the behavioral epoch to which it is being compared, *D_2_*, each row corresponds to timepoints and each column to neurons. For each matrix, we then compute matrices *P_1_* and *P_2_* that are composed of 6 principal component vectors as columns. The alignment index, *a,* is then computed using

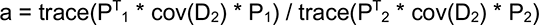

In each case, the alignment index was computed again after reversing the order of the behavioral epochs, and the mean of the resulting values was reported.

#### Two-photon Imaging Analysis

GCaMP6f fluorescence image time series were corrected for motion artifacts offline using nonrigid alignment via the normcorre algorithm (Pnevmatikakis and Giovannucci, 2017). Individual dendritic cross-sections (sources) were defined from motion-corrected image time series using the CNMF function library (Giovannucci et al., 2019). Input to the ‘initialize_components’ function was defined using the watershed algorithm (MATLAB function ‘watershed’) on the mean time series image. The output of ‘initialize_components’ was further refined using the ‘manually_refine_components’ function. Sources were then further refined using the ‘update_spatial_components’ function (‘spatial_method’ = ‘constrained’, ‘search_method’ = ‘ellipse’, ‘min_size’ = 1, ‘max_size’ = 5). To remove slow shifts in fluorescence signal baseline and improve subsequent spike inference, fluorescence time series for the resulting sources were then baseline corrected by identifying the tenth percentile value from each successive 25-sample time series segment and subtracting this value from samples within the given segment. Calcium fluctuations were then estimated from source fluorescence signals and spiking activity was inferred from estimated calcium fluctuations using the ‘cont_ca_sampler’ and ‘make_mean_sample’ functions (‘deconv_method’ = ‘MCMC’).

To select trial equivalents from self-paced behavior, segments of EMG time series were identified from the alternation 1, cocontraction, and alternation 2 epochs, as well as from epochs of muscle quiescence, using methods similar to those described above for analysis of electrical recordings. Estimated calcium and spike time series for each source were aligned to EMG time series using the timing of the collection of the image line closest to the centroid of the source in each frame. Estimated calcium and spike time series for each source were smoothed with a Gaussian having a 10 ms standard deviation. For each source, segments of estimated calcium and spike time series corresponding to identified EMG time series segments for each behavioral epoch were assembled to generate estimated calcium and spiking trial-averages.

Most sources demonstrated little detectable activity or little behaviorally correlated activity. Because the total number of sources was large and subsequent identification of rabies-labeled sources involved manual intervention, sources lacking relevant activity were eliminated from further analysis using the following method. The goal of this method was to eliminate sources that did not have a substantial projection onto any of the top principal components computed for the trial-averaged estimated calcium of sources identified in a given image time series. Principal component analysis was performed on the S sources by T time points matrix formed by the estimated calcium trial averages for the alternation 1 and cocontraction epochs during a given image time series. We computed the distribution of the absolute values of the projections of all sources onto the half of principal components explaining the least data variance (i.e., the second half of components when ordered by variance explained) and computed the mean and standard deviation of this distribution. Sources were eliminated unless they had a projection onto one of the top 7 principal components for either the alternation 1 or cocontraction epoch that was larger than 6 standard deviations above the mean of this distribution of projections onto the second half of components.

Sources from each GCaMP6f image time series were then classified as either tdTomato positive (rabies-labeled), tdTomato negative, or undetermined using the following method. We calculated the two-dimensional correlation (MATLAB function ‘corr2’) between the mean of the GCaMP6f image time series and each GCaMP6f image from the Z stack collected following rabies injection, after using nonrigid motion correction via the normcorre algorithm to align the two images. The GCaMP6f Z stack image yielding the highest correlation was selected for further analysis, along with the corresponding tdTomato image collected in the same plane. Any offset between the selected GCaMP6f and tdTomato Z stack images was then calculated using the normalized two-dimensional cross-correlation (‘findoff’ function downloaded from the MATLAB File Exchange on 11/1/2017) and corrected by shifting one image relative to the other. The pixel masks for each source were then mapped to the selected GCaMP6f Z stack image using the ‘shift_reconstruct’ function from the CNMF library. Outlines of pixel masks were defined by thresholding masks at 25% of the maximum mask value, taking the largest connected component among the suprathreshold pixels, and taking the outline of that component. These outlines were then overlaid onto the selected, aligned GCaMP6f and tdTomato Z stack images for visual inspection and source classification.

Sources were classified as tdTomato positive if (a) one cross-section in the GCaMP6f Z stack image could be unambiguously linked to the source mask, and (b) that cross-section showed obvious tdTomato labeling above background. Sources were classified as tdTomato negative if (a) one cross-section in the GCaMP6f Z stack image could be unambiguously linked to the source mask, but (b) that cross-section showed no discernable tdTomato labeling above background. The remaining sources were left unclassified. These undetermined sources were a majority of the overall number of sources meeting the above PCA-based selection criteria. In many cases, source outlines did not overlap with a GCaMP6f Z stack cross-section or overlap one completely, likely owing to micron-scale movement of dendrites relative to one another between the GCaMP6f image time series collection and Z stack collection after rabies injection (10-15 days). In a minority of these cases, source outlines could still be unambiguously linked to certain cross-sections based on the relative arrangement of nearby cross-sections. In many other cases, cross-sections in the GCaMP6f Z stack image could not be unambiguously classified as tdTomato positive or negative because of ambiguity as to whether tdTomato signal was above background or ambiguity about whether apparent tdTomato signal had come from a neighboring cross-section.

Since GCaMP6f fluorescence-based spike inference will deviate from actual underlying spiking in a manner that could exaggerate the appearance of task-specific neurons, we sought to account for this effect in comparing specificity indices for optically recorded corticospinal neurons and electrically recorded layer 5b neurons. For this analysis, we began with our spike times for electrically recorded wide-waveform layer 5b neurons that met the inclusion requirements for the analysis of firing described above. We then used a procedure to simulate GCaMP6f fluorescence-based inferred spiking for each of these neurons. The procedure involved two free parameters: a base transient height and a noise scale factor. The same parameter values were used for all neurons, but their values were set so as to match in the results the ratio between the mean detected spike rate for our optically recorded corticospinal neurons and electrically recorded wide-waveform layer 5b neurons, ∼1/6. That is, parameters were set so that the resulting inferred spike rate from simulated fluorescence was on average 1/6 of the original mean firing rate across neurons.

The simulation procedure went as follows. For each neuron, a time series vector was generated that was zero everywhere except at spike times, where the magnitude was a constant chosen at random from a uniform distribution ranging from 0.2x to 1.8x the base transient height parameter. Each nonzero magnitude in the vector was then randomly varied to match observed variation in single spike calcium transient heights (Éltes et al., 2019). Vectors were then convolved with a double exponential kernel with rise and fall time constants matching observed values for GCaMP6f transients. The resulting vectors were then transformed again using the Hill equation, assuming published values for the Hill coefficient (2.27) and K_d_ (375 nM; Rose et al., 2014). Finally, Gaussian noise was added that had a size equal to the noise scale factor times the transient magnitude constant chosen for the given neuron, producing a time series simulating a GCaMP6f fluorescent measurement. Spiking was then inferred from these simulated fluorescent measurements as above using the ‘cont_ca_sampler’ and ‘make_mean_sample’ functions (Giovannucci et al., 2019).

---

**Figure S1.**
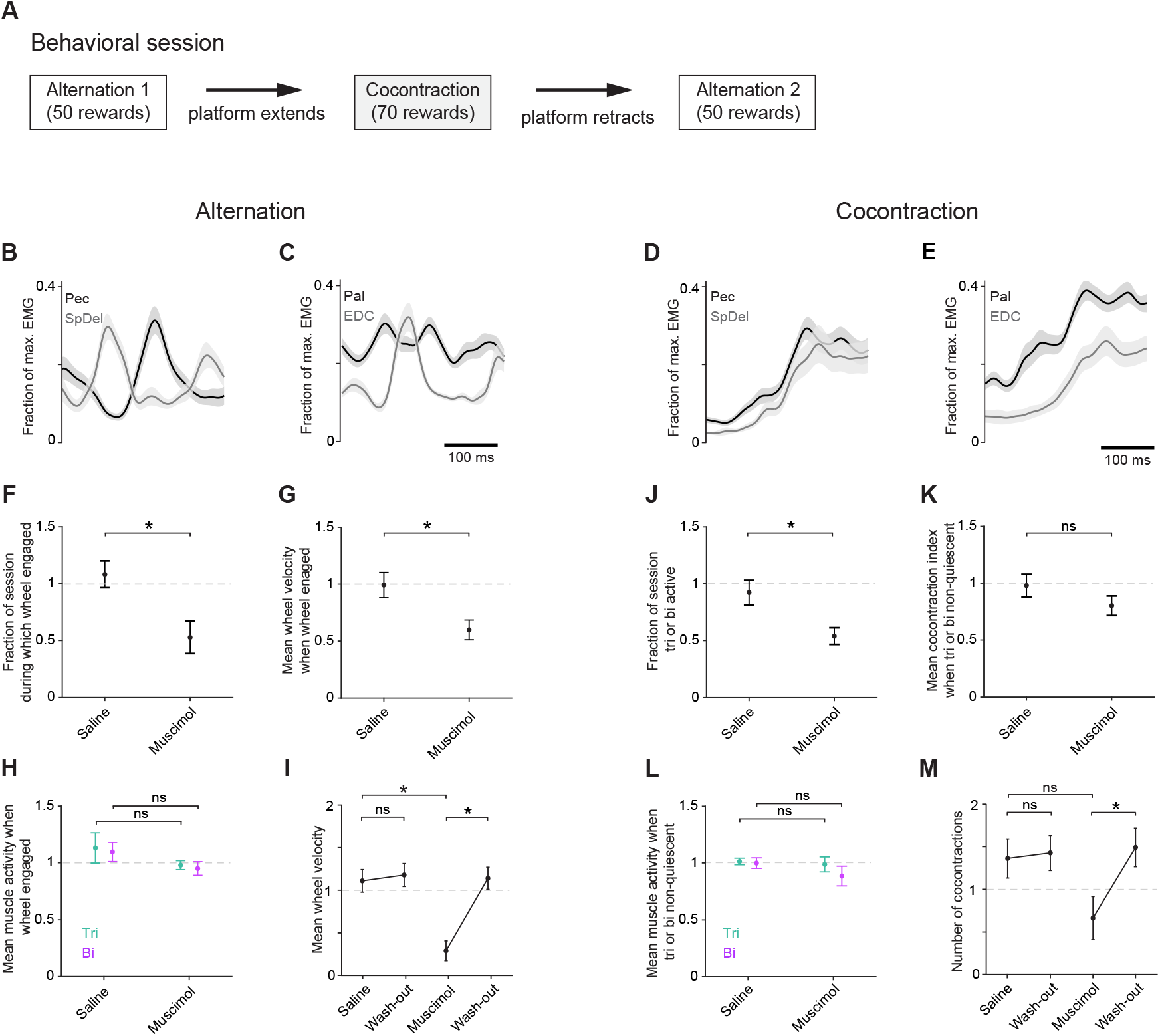
Additional EMG and effects of muscimol on behavior. (A) Flow chart depicting the order of tasks in the behavioral paradigm. (B-E) Trial-averaged EMG ± SEM (n = 30 trials) aligned to triceps onset for pectoralis (Pec, black) and spinodeltoidus (SpDel, gray) during alternation (B) and cocontraction (D) and for palmaris longus (Pal, black) and extensor digitorum communis (EDC, gray) during alternation (C) and cocontraction (E). B-C and D-E show data from the same time points as shown in Figure 1 F-G and J-K, respectively. Panels F-M show mean ± SEM (n = 5 mice) of behavioral metrics 90 minutes after injection of muscimol or saline to the CFA, assessed over the first 20 minutes of behavioral session and normalized to baseline (performance averaged over the three sessions preceding injection). (F) Fraction of session during which the wheel was engaged (moving towards the mouse) during alternation (paired one-tailed t-test p=0.021). (G) Mean wheel velocity only during times the wheel was engaged (p=0.049), (H) Mean muscle activity during times the wheel was engaged (Tri p = 0.129; Bi p = 0.138). (I) Mean wheel velocity after bilateral injection of saline p=0.225 or muscimol p=0.003. (J) Fraction of session during which either triceps or biceps was active (> 10% maximum contraction) during cocontraction (p=0.013). (K) Mean cocontraction index during times when either triceps or biceps was active (p=0.173). (L) Mean muscle activity during times when either muscle was active (Tri p=0.357, Bi p=0.206). (M) Number of cocontraction events after bilateral injection of saline (p=0.350) or muscimol (p=0.028).

**Figure S2.**
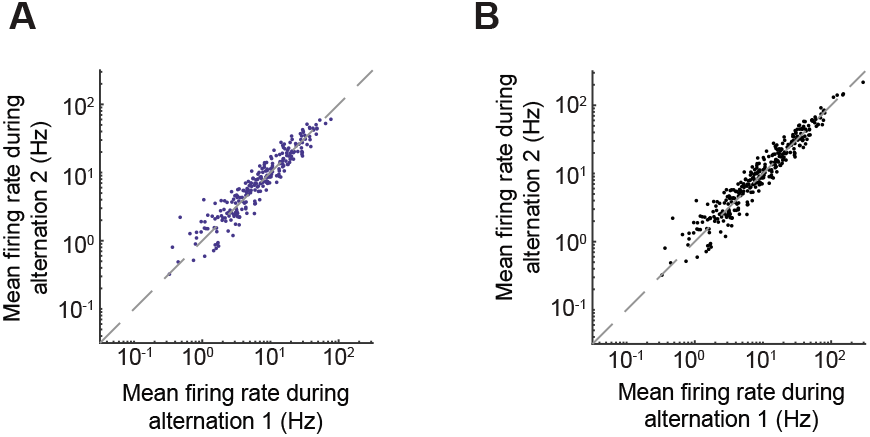
All neurons demonstrate a continuum of task specificity. (A-B) The mean firing rate during the first epoch of alternation (the higher rate when comparing flexion and extension was chosen) plotted against the mean firing rate during the second epoch of alternation for neurons that were included for analysis among wide-waveform (A, blue) and all neurons (B, black). Neurons whose mean firing rates were dramatically different during these epochs were excluded from analysis.

**Figure S3.**
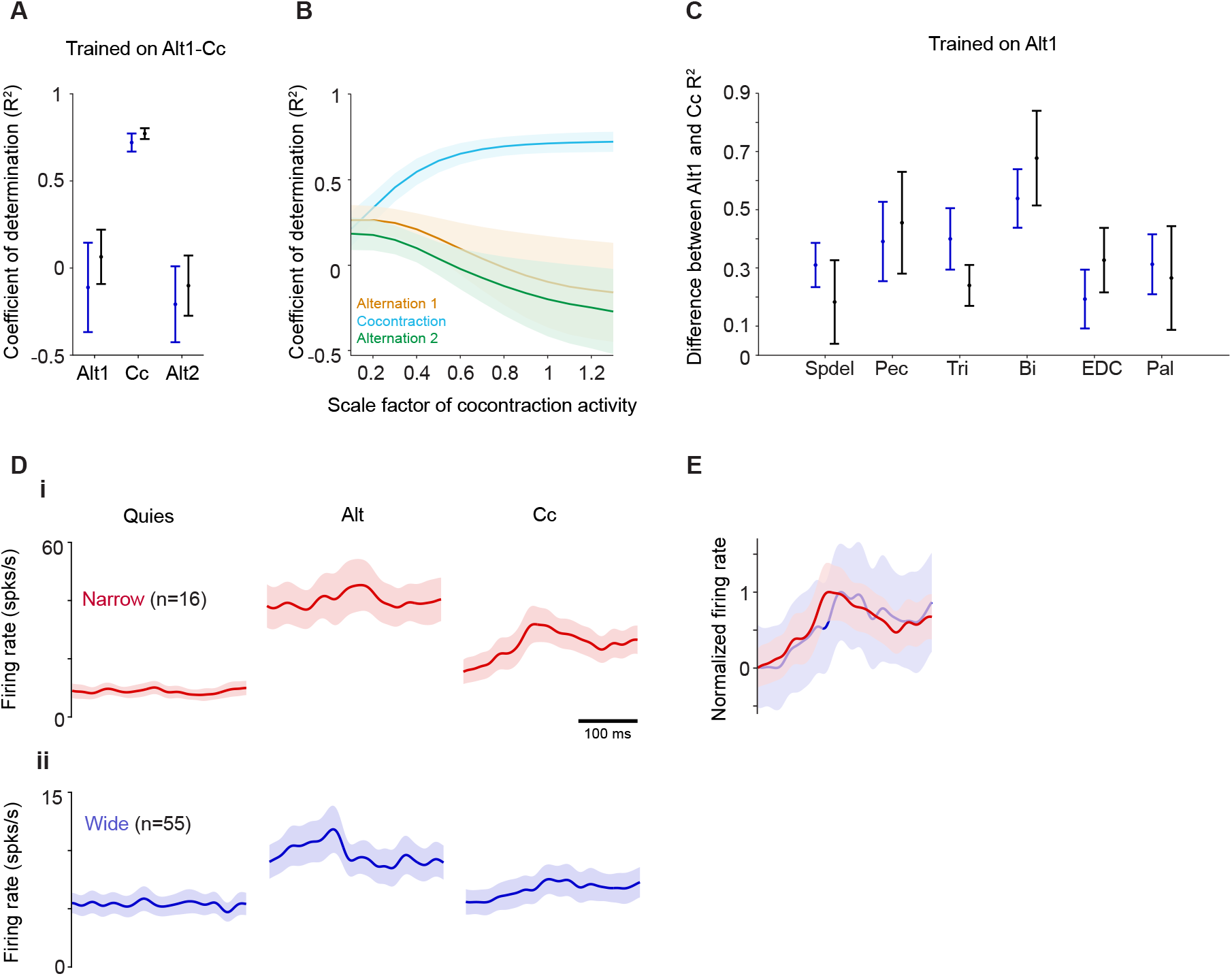
Additional support for neural activity task-specific rather than cocontraction the sum of flexion and extension. (A) Mean ± SEM (n=4 mice) of the coefficient of determination (R^2^) for wide-waveform neurons (blue) and all neurons (black) for activity during the first epoch of alternation (Alt1), cocontraction (Cc), and the second epoch of alternation (Alt1) for a model trained on trials from both alternation 1 and cocontraction concatenated in time. (B) Mean ± SEM (n=4 mice) of the coefficient of determination for wide-waveform neurons when the cocontraction neural and activity across the six recorded muscles is scaled by a factor from 0.1-1.3 for trials from alternation 1 (orange), cocontraction (blue), and alternation 2 (green). (C) Mean ± SEM (n=4 mice) of the differences between the R^2^ values during alternation 1 and cocontraction for the model trained on alternation 1 wide-waveform (blue) and all neurons (black) activity showing that no one muscle drives the failure in generalization. (D) Example trial-average activity ± SEM during muscle quiescence, alternation, and cocontraction of narrow-waveform neurons (i; n=16 neurons) and of wide-waveform neurons (ii; n=55 neurons) from a single mouse. (E) Example of trial-average activity ± SEM of narrow- (red) and wide- (blue) waveform neurons normalized to respective minima and maxima from a single mouse.

**Figure S4.**
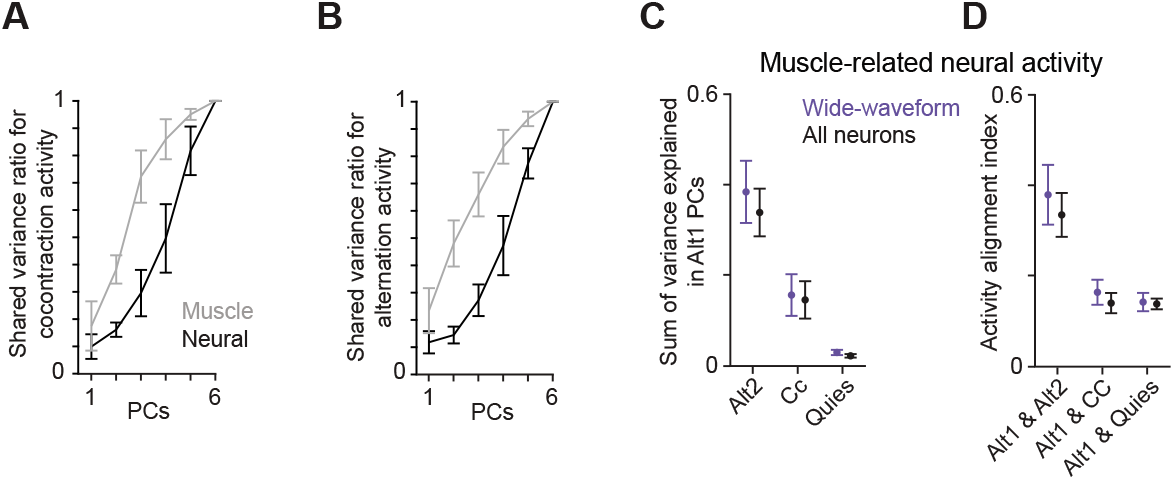
Task-specific motor cortical activity covariation controls. (A) Shared variance among muscle (gray) and all neural (black) activity for the variance of cocontraction activity in the top alternation 1 PCs relative to the variance of cocontraction in its own top PCs. (B) Shared variance of alternation 1 activity in the top cocontraction PCs relative to the variance of alternation 1 in its own top PCs. (C-D) For only the muscle-related activity extracted from all neurons (black) and wide-waveform neurons (blue), mean ± SEM (n=4 mice) of the sum of variance explained in the top 6 alternation 1 PCs (C) and of the alignment of firing rates in alternation 1 with other epochs (D).

**Figure S5.**
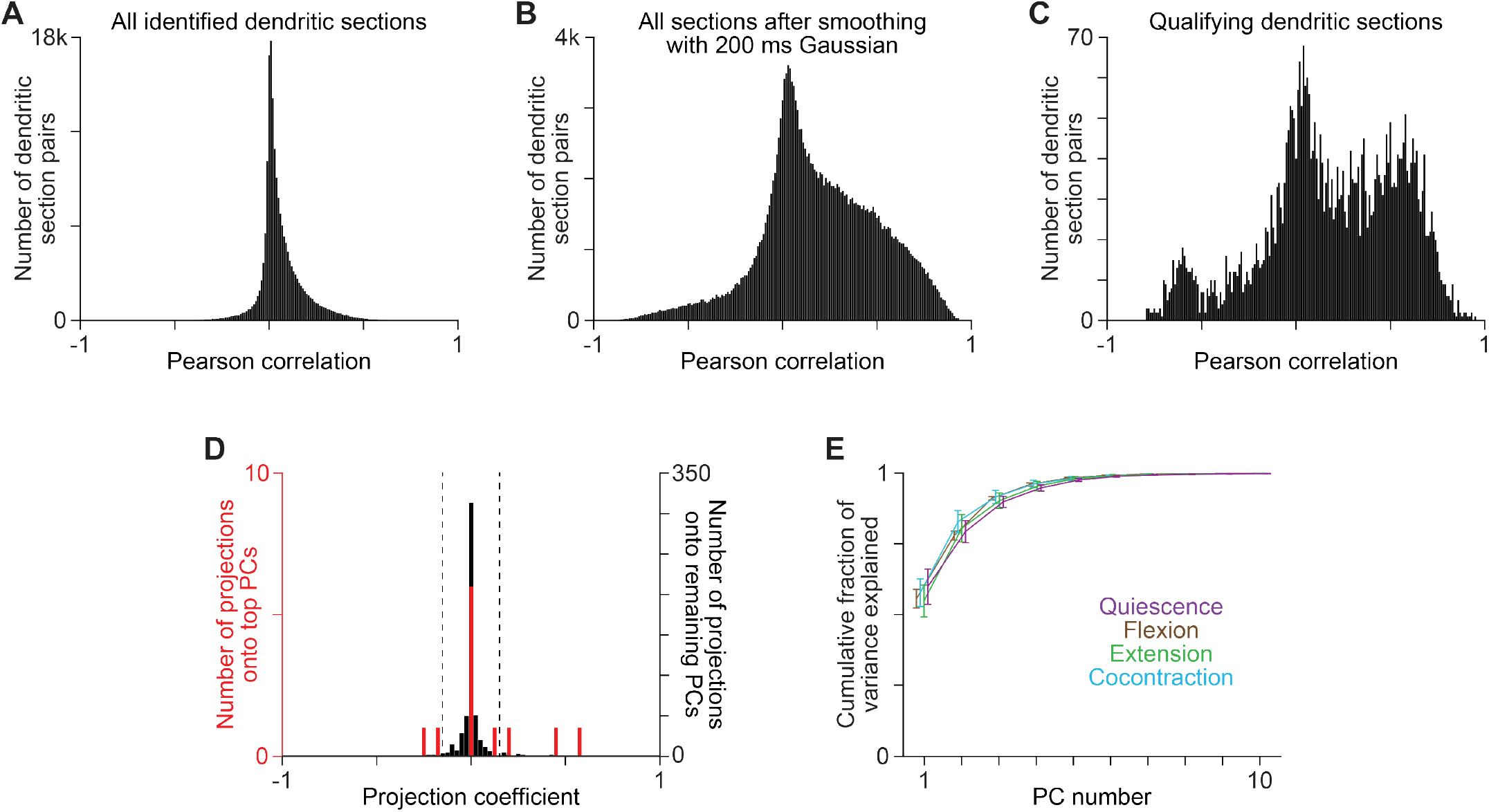
GCaMP6f fluorescence measurements. (A-C) Histograms of Pearson correlation for all possible pairs of dendritic fluorescence time series recorded in individual planes. These plots did not reveal a small subset of pairs with distinctly high correlations suggestive of dendrites emanating from a common soma, not for all pairs (A), all pairs after smoothing fluorescence time series with a 200 ms Gaussian (B), nor for pairs comprised of only qualifying dendritic sections (C). (D) Histograms of the projection of an example neuron onto all principal components for dendritic sections measured in one imaging window. Principal components were computed separately for alternation and cocontraction and the projections aggregated. Histograms are separated for projections onto the top 6 components from either behavioral epoch (red) or all remaining components (black). Dotted lines show the mean ± three standard deviations of the latter distribution of projections. Dendritic sections qualified for subsequent analysis if at least one projection onto a top 6 component fell outside these boundaries, indicating a significant degree of signal in the time series for that section. (E) Cumulative variance explained by the top 10 principal components for qualifying dendritic sections for four behavioral epochs. In all cases, 6 principal components accounted for ∼99% of activity variance.

## References

Aimonetti, J.-M.M., and Nielsen, J.B. (2002). Cortical excitability and motor task in man: an investigation of the wrist extensor motor area. Experimental Brain Research 143, 431–439.

Akay, T., Acharya, H.J., Fouad, K., and Pearson, K.G. (2006). Behavioral and Electromyographic Characterization of Mice Lacking EphA4 Receptors. J Neurophysiol 96, 642– 651.

Albert, S.T., Hadjiosif, A.M., Jang, J., Zimnik, A.J., Soteropoulus, D.S., Baker, S.N., Churchland, M.M., Krakauer, J.W., and Shadmehr, R. (2020). Postural control of arm and fingers through integration of movement commands. eLife 9:e52507.

Ames, K., and Churchland, M.M. (2019). Motor cortex signals for each arm are mixed across hemispheres and neurons yet partitioned within the population response. eLife 8, e46159.

Ayling, O.G., Harrison, T.C., Boyd, J.D., Goroshkov, A., and Murphy, T.H. (2009). Automated light-based mapping of motor cortex by photoactivation of channelrhodopsin-2 transgenic mice. Nature Methods 6, 219–224.

Azim, E., Jiang, J., Alstermark, B., and Jessell, T.M. (2014). Skilled reaching relies on a V2a propriospinal internal copy circuit. Nature 508, 357–363.

Barthó, P., Hirase, H., Monconduit, L., Zugaro, M., Harris, K.D., and Buzsáki, G. (2004). Characterization of Neocortical Principal Cells and Interneurons by Network Interactions and Extracellular Features. Journal of Neurophysiology 92, 600–608.

Beaulieu, C. (1993). Numerical data on neocortical neurons in adult rat, with special reference to the GABA population. Brain Res 609, 284–292.

Brown, A.R., and Teskey, C.G. (2014). Motor Cortex Is Functionally Organized as a Set of Spatially Distinct Representations for Complex Movements. The Journal of Neuroscience 34, 13574–13585.

Capaday, C. (2004). The integrated nature of motor cortical function. The Neuroscientist : A Review Journal Bringing Neurobiology, Neurology and Psychiatry 10, 207–220.

Capaday, C., Devanne, H., Bertrand, L., and Lavoie (1998). Intracortical connections between motor cortical zones controlling antagonistic muscles in the cat: a combined anatomical and physiological study. Experimental Brain Research 120, 223–232.

Capaday, C., Ethier, C., Vreeswijk, C., and Darling, W.G. (2013). On the functional organization and operational principles of the motor cortex. Frontiers in Neural Circuits 7, 66.

Chen, X.Y., Chen, L., Chen, Y., and Wolpaw, J.R. (2006a). Operant conditioning of reciprocal inhibition in rat soleus muscle. Journal of Neurophysiology 96, 2144–2150.

Chen, X.Y., Chen, Y., Chen, L., Tennissen, A.M., and Wolpaw, J.R. (2006b). Corticospinal tract transection permanently abolishes H-reflex down-conditioning in rats. Journal of Neurotrauma 23, 1705–1712.

Churchland, M.M., and Shenoy, K.V. (2007). Temporal Complexity and Heterogeneity of Single-Neuron Activity in Premotor and Motor Cortex. J Neurophysiol 97, 4235–4257.

Churchland, M.M., Cunningham, J.P., Kaufman, M.T., Foster, J.D., Nuyujukian, P., Ryu, S.I., and Shenoy, K.V. (2012). Neural population dynamics during reaching. Nature 487, 51–56.

Daie, K., Goldman, M.S., and Aksay, E. (2015). Spatial Patterns of Persistent Neural Activity Vary with the Behavioral Context of Short-Term Memory. Neuron 85, 847–860.

Dombeck, D., Graziano, and Tank, D. (2009). Functional Clustering of Neurons in Motor Cortex Determined by Cellular Resolution Imaging in Awake Behaving Mice. Journal of Neuroscience 29, 13751–13760.

Druckmann, S., and Chklovskii, D.B. (2012). Neuronal circuits underlying persistent representations despite time varying activity. Current Biology: CB 22, 2095–2103.

Eccles, R., and Lundberg, A. (1958). Integrative pattern of Ia synaptic actions on motoneurones of hip and knee muscles. J Physiol. 144(*2*), 271–298.

Elsayed, G.F., Lara, A.H., Kaufman, M.T., Churchland, M.M., and Cunningham, J.P. (2016). Reorganization between preparatory and movement population responses in motor cortex. Nat Commun 7, 13239.

Éltes, T., Szoboszlay, M., Kerti-Szigeti, K., Nusser, Z. (2019). Improved spike inference accuracy by estimating the peak amplitude of unitary [Ca2+] transients in weakly GCaMP6f-expressing hippocampal pyramidal cells. J Physiol 597, 2925–2947.

Estebanez, L., Hoffmann, D., Voigt, B.C., and Poulet, J. (2017). Parvalbumin-Expressing GABAergic Neurons in Primary Motor Cortex Signal Reaching. Cell Reports 20, 308–318.

Ethier, C., Brizzi, L., Giguère, D., and Capaday, C. (2007). Corticospinal control of antagonistic muscles in the cat. The European Journal of Neuroscience 26, 1632–1641.

Feldman, A.G. (1980). Superposition of motor programs—II. Rapid forearm flexion in man. Neuroscience 5, 91–95.

Fetz, E., and Cheney, P. (1987). Functional relations between primate motor cortex cells and muscles: fixed and flexible. Ciba Foundation Symposium 132, 98–117.

Fink, A.J., Croce, K.R., Huang, J.Z., Abbott, L., Jessell, T.M., and Azim, E. (2014). Presynaptic inhibition of spinal sensory feedback ensures smooth movement. Nature 509, 43–48.

Fritsch, G., and Hitzig, E. (1870). Electric excitability of the excitability of the cerebrum (Uber die elektrische Erregbarkeit des Grosshirns). Epilepsy Behav 15, 123–130.

Gallego, J.A., Perich, M.G., Miller, L.E., and Solla, S.A. (2017). Neural Manifolds for the Control of Movement. Neuron 94, 978–984.

Giovannucci, A., Friedrich, J., Gunn, P., Kalfon, J., Brown, B.L., Koay, S., Taxidis, J., Najafi, F., Gauthier, J.L., Zhou, P., et al. (2019). CaImAn an open source tool for scalable calcium imaging data analysis. Elife 8, e38173.

Graziano, M., Taylor, C., and Moore, T. (2002). Complex Movements Evoked by Microstimulation of Precentral Cortex. Neuron 34, 841–851.

Graziano, M.S., Aflalo, T.N., and Cooke, D.F. (2005). Arm Movements Evoked by Electrical Stimulation in the Motor Cortex of Monkeys. J Neurophysiol 94, 4209–4223.

Graziano, M.S. (2006). Progress in understanding spatial coordinate systems in the primate brain. Neuron. 51(1):7–9.

Gribble, P.L., Mullin, L.I., Cothros, N., and Mattar, A. (2003). Role of Cocontraction in Arm Movement Accuracy. Journal of Neurophysiology 89, 2396–2405.

Greenberg, D.S., Wallace, D.J., Voit, K.-M., Wuertenberger, S., Czubayko, U., Monsees, A., Handa, T., Vogelstein, J.T., Seifert, R., Groemping, Y., Kerr, J.N.D. (2021). Accurate action potential inference from a calcium sensor protein through biophysical modeling. bioRxiv 2018.11.29.479055.

Guo, Z.V., Li, N., Huber, D., Ophir, E., Gutnisky, D., Ting, J.T., Feng, G., and Svoboda, K. (2014a). Flow of cortical activity underlying a tactile decision in mice. Neuron 81, 179–194.

Guo, Z.V., Hires, S., Li, N., O’Connor, D.H., Komiyama, T., Ophir, E., Huber, D., Bonardi, C., Morandell, K., Gutnisky, D., et al. (2014b). Procedures for behavioral experiments in head-fixed mice. PloS One 9, e88678.

Harrison, T.C., Ayling, O.G., and Murphy, T.H. (2012). Distinct cortical circuit mechanisms for complex forelimb movement and motor map topography. Neuron 74, 397–409.

Huang, H.J., Kram, R., and Ahmed, A.A. (2012). Reduction of Metabolic Cost during Motor Learning of Arm Reaching Dynamics. The Journal of Neuroscience 32, 2182–2190.

Humphrey, D.R., and Reed, D.J. (1983). Separate cortical systems for control of joint movement and joint stiffness: Reciprocal activation and coactivation of antagonist muscles. Motor Control Mechanisms in Health and Disease 347–372.

Isomura, Y., Harukuni, R., Takekawa, T., Aizawa, H., and Fukai, T. (2009). Microcircuitry coordination of cortical motor information in self-initiation of voluntary movements. Nature Neuroscience 12, 1586–1593.

Jankowska, E. (1992). Interneuronal relay in spinal pathways from proprioceptors. Progress in Neurobiology 38, 335–378.

Jankowska, E., Padel, Y., and Tanaka, R. (1976). Disynaptic inhibition of spinal motoneurones from the motor cortex in the monkey. The Journal of Physiology 258(*2*), 467–487.

Johannsen, P., Christensen, L., Sinkjær, T., and Nielsen, J. (2001). Cerebral functional anatomy of voluntary contractions of ankle muscles in man. The Journal of Physiology 535, 397–406.

Kargo, W.J., and Nitz, D.A. (2004). Improvements in the Signal-to-Noise Ratio of Motor Cortex Cells Distinguish Early versus Late Phases of Motor Skill Learning. The Journal of Neuroscience 24, 5560–5569.

Kaufman, M.T., Churchland, M.M., and Shenoy, K.V. (2013). The roles of monkey M1 neuron classes in movement preparation and execution. Journal of Neurophysiology 110, 817–825.

Kaufman, M.T., Churchland, M.M., Ryu, S.I., and Shenoy, K.V. (2014). Cortical activity in the null space: permitting preparation without movement. Nature Neuroscience 17, 440–448.

Knarr, B.A., Zeni, J.A., and Higginson, J.S. (2012). Comparison of electromyography and joint moment as indicators of co-contraction. Journal of Electromyography and Kinesiology 22, 607– 611.

Komiyama, T., Sato, T.R., O’Connor, D.H., Zhang, Y.-X.X., Huber, D., Hooks, B.M., Gabitto, M., and Svoboda, K. (2010). Learning-related fine-scale specificity imaged in motor cortex circuits of behaving mice. Nature 464, 1182–1186.

Kvitsiani, D., Ranade, S., Hangya, B., Taniguchi, H., Huang, J., and Kepecs, A. (2013). Distinct behavioural and network correlates of two interneuron types in prefrontal cortex. Nature 498, 363.

Lacquaniti, F., and Maioli, C. (1987). Anticipatory and reflex coactivation of antagonist muscles in catching. Brain Research 406, 373–378.

Leutgeb, S., Leutgeb, J.K., Barnes, C.A., Moser, E.I., McNaughton, B.L., and Moser, M.-B. (2005). Independent Codes for Spatial and Episodic Memory in Hippocampal Neuronal Ensembles. Science 309(5734), 619–23.

Lévénez, M., Garland, S., Klass, M., and Duchateau, J. (2008). Cortical and spinal modulation of antagonist coactivation during a submaximal fatiguing contraction in humans. Journal of Neurophysiology 99, 554–563.

Leyton, A., and Sherrington, C. (1917). Observations on the Excitable Cortex of the Chimpanzee, Orang-utan, and Gorilla. Q J Exp Physiol 11, 135–222.

Mante, V., Sussillo, D., Shenoy, K.V., and Newsome, W.T. (2013). Context-dependent computation by recurrent dynamics in prefrontal cortex. Nature 503, 78–84.

Matsumura, M., Sawaguchi, T., Oishi, T., Ueki, K., and Kubota, K. (1991). Behavioral deficits induced by local injection of bicuculline and muscimol into the primate motor and premotor cortex. Journal of Neurophysiology 65, 1542–1553.

Mazzocchio, R., Rossi, A., and Rothwell, J. (1994). Depression of Renshaw recurrent inhibition by activation of corticospinal fibres in human upper and lower limb. The Journal of Physiology 481 (*Pt 2*), 487–498.

McCormick, D., Connors, B., Lighthall, J., and Prince, D. (1985). Comparative electrophysiology of pyramidal and sparsely spiny stellate neurons of the neocortex. J Neurophysiol 54, 782–806.

Milner, T.E. (2002). Adaptation to destabilizing dynamics by means of muscle cocontraction. Experimental Brain Research 143, 406–416.

Miri, A., Warriner, C.L., Seely, J.S., Elsayed, G.F., Cunningham, J.P., Churchland, M.M., and Jessell, T.M. (2017). Behaviorally Selective Engagement of Short-Latency Effector Pathways by Motor Cortex. Neuron 95, 683–696.e11.

Mittmann, W., Wallace, D.J., Czubayko, U., Herb, J.T., Schaefer, A.T., Looger, L.L., Denk, W., and Kerr, J.N. (2011). Two-photon calcium imaging of evoked activity from L5 somatosensory neurons in vivo. Nat Neurosci 14, 1089–1093.

Nielsen, J., and Kagamihara, Y. (1992). The regulation of disynaptic reciprocal Ia inhibition during co-contraction of antagonistic muscles in man. The Journal of Physiology 456, 373–391.

Nielsen, J., and Kagamihara, Y. (1993). The regulation of presynaptic inhibition during co-contraction of antagonistic muscles in man. The Journal of Physiology 464, 575–593.

Nielsen, J., Petersen, N., Deuschl, G., and Ballegaard, M. (1993). Task-related changes in the effect of magnetic brain stimulation on spinal neurones in man. The Journal of Physiology 471, 223–243.

Pandarinath, C., Ames, K., Russo, A.A., Farshchian, A., Miller, L.E., Dyer, E.L., and Kao, J.C. (2018). Latent Factors and Dynamics in Motor Cortex and Their Application to Brain-Machine Interfaces. The Journal of Neuroscience: The Official Journal of the Society for Neuroscience 38, 9390–9401.

Pearson, K.G., Acharya, H., and Fouad, K. (2005). A new electrode configuration for recording electromyographic activity in behaving mice. J Neurosci Meth 148, 36–42.

Penfield, W., and Rasmussen, T. (1950). The cerebral cortex of man: a clinical study of localization of function. JAMA 144(60), 1412.

Perez, M.A., Lundbye-Jensen, J., and Nielsen, J.B. (2007). Task-specific depression of the soleus H-reflex after cocontraction training of antagonistic ankle muscles. Journal of Neurophysiology 98, 3677–3687.

Perich, M.G., Conti, S., Badi, M., Bogaard, A., Barra, B., Wurth, S.M., Bloch, J., Courtine, G., Micera, S., Capogrosso, M., Milevovic, T. (2020). Motor cortical dynamics are shaped by multiple distinct subspaces during naturalistic behavior. bioRxiv 2020.07.30.228767

Perich, M.G., Gallego, J.A., and Miller, L.E. (2018). A Neural Population Mechanism for Rapid Learning. Neuron 100, 964–976.e7.

Peters, A.J., Lee, J., Hedrick, N.G., O’Neil, K., and Komiyama, T. (2017). Reorganization of corticospinal output during motor learning. Nat Neurosci 20, 1133–1141.

Pho, G.N., Goard, M.J., Woodson, J., Crawford, B., and Sur, M. (2018). Task-dependent representations of stimulus and choice in mouse parietal cortex. Nat Commun 9, 2596.

Pinto, L., Rajan, K., DePasquale, B., Thiberge, S.Y., Tank, D.W., and Brody, C.D. (2019). Task-Dependent Changes in the Large-Scale Dynamics and Necessity of Cortical Regions. Neuron.

Pnevmatikakis, E.A., and Giovannucci, A. (2017). NoRMCorre: An online algorithm for piecewise rigid motion correction of calcium imaging data. J Neurosci Meth 291, 83–94.

Rathelot, J.-A.A., and Strick, P.L. (2009). Subdivisions of primary motor cortex based on cortico-motoneuronal cells. Proceedings of the National Academy of Sciences of the United States of America 106, 918–923.

Reardon, T.R., Murray, A.J., Turi, G.F., Wirblich, C., Croce, K.R., Schnell, M.J., Jessell, T.M., and Losonczy, A. (2016). Rabies Virus CVS-N2c(ΔG) Strain Enhances Retrograde Synaptic Transfer and Neuronal Viability. Neuron 89, 711–724.

Rose, T., Goltstein, P.M., Portugues, R., Griesbeck, O. (2014). Putting a finishing touch on GECIs. Front Mol Neurosci. 7:88.

Rossant, C., Kadir, S., Goodman, D., and Schulman, J. (2016). Spike sorting for large, dense electrode arrays. Nature Neuroscience 19, 634–641.

Rudolph, K.S., Axe, M.J., and Snyder-Mackler, L. (2000). Dynamic stability after ACL injury: who can hop? Knee Surgery, Sports Traumatology, Arthroscopy 8, 262–269.

Russ, J.B., Verina, T., Comer, J.D., Comi, A.M., and Kaltschmidt, J.A. (2013). Corticospinal tract insult alters GABAergic circuitry in the mammalian spinal cord. Frontiers in Neural Circuits 7, 150.

Russo, A.A., Bittner, S.R., Perkins, S.M., Seely, J.S., London, B.M., Lara, A.H., Miri, A., Marshall, N.J., Kohn, A., Jessell, T.M., et al. (2018). Motor Cortex Embeds Muscle-like Commands in an Untangled Population Response. Neuron 97, 953–966.e8.

Sadtler, P.T., Quick, K.M., Golub, M.D., Chase, S.M., Ryu, S.I., Tyler-Kabara, E.C., Yu, B.M., and Batista, A.P. (2014). Neural constraints on learning. Nature 512, 423.

Saliba, C.M., Rainbow, M.J., Selbie, W.S., Deluzio, K.J., Scott, S.H. (2020). Co-contraction uses dual-control of agonist-antagonist muscles to improve motor performance. BioRxiv.

Scheidt, R.A., and Ghez, C. (2007). Separate Adaptive Mechanisms for Controlling Trajectory and Final Position in Reaching. J Neurophysiol 98, 3600–3613.

Schneider, C., Devanne, H., Lavoie, B.A., and Capaday, C. (2002). Neural mechanisms involved in the functional linking of motor cortical points. Experimental Brain Research 146, 86– 94.

Schroeder, K.E., Perkins, S.M., Wang, Q., Churchland, M.M. (2021). Cortical control of virtual self-motion using task-specific subspaces. Journal of Neuroscience 42(2), 220–239.

Semedo, J.D., Zandvakili, A., Machens, C.K., Yu, B.M., and Kohn, A. (2019). Cortical Areas Interact through a Communication Subspace. Neuron 102, 249–259.e4.

Shadmehr, R. (2017). Distinct neural circuits for control of movement vs. holding still. Journal of Neurophysiology 117, 1431–1460.

Smith, A.M. (1981). The coactivation of antagonist muscles. Can J Physiol Pharm 59, 733–747.

Sussillo, D., Churchland, M.M., Kaufman, M.T., and Shenoy, K.V. (2015). A neural network that finds a naturalistic solution for the production of muscle activity. Nat Neurosci 18, 1025–1033.

Taniguchi, H., He, M., Wu, P., Kim, S., Paik, R., Sugino, K., Kvitsiani, D., Kvitsani, D., Fu, Y., Lu, J., et al. (2011). A resource of Cre driver lines for genetic targeting of GABAergic neurons in cerebral cortex. Neuron 71, 995–1013.

Tennant, K.A., Adkins, D.L., Donlan, N.A., Asay, A.L., Thomas, N., Kleim, J.A., and Jones, T.A. (2011). The Organization of the Forelimb Representation of the C57BL/6 Mouse Motor Cortex as Defined by Intracortical Microstimulation and Cytoarchitecture. Cerebral Cortex 21, 865–876.

Tervo, G.R.D., Hwang, B.-Y., Viswanathan, S., Gaj, T., Lavzin, M., Ritola, K.D., Lindo, S., Michael, S., Kuleshova, E., Ojala, D., et al. (2016). A Designer AAV Variant Permits Efficient Retrograde Access to Projection Neurons. Neuron 92, 372–382.

Thoroughman, K.A., and Shadmehr, R. (1999). Electromyographic Correlates of Learning an Internal Model of Reaching Movements. Journal of Neuroscience 19, 8573–8588.

Tilney, F., and Pike, F.H. (1925). Muscular Coordination Experimentally Studied in its Relation to the Cerebellum. Archives Neurology Psychiatry 13, 289–334.

Watabe-Uchida, M., Zhu, L., Ogawa, S.K., Vamanrao, A., and Uchida, N. (2012). Whole-Brain Mapping of Direct Inputs to Midbrain Dopamine Neurons. Neuron 74, 858–873.

Zhao, S., Ting, J.T., Atallah, H.E., Qiu, L., Tan, J., Gloss, B., Augustine, G.J., Deisseroth, K., Luo, M., Graybiel, A.M., et al. (2011). Cell type–specific channelrhodopsin-2 transgenic mice for optogenetic dissection of neural circuitry function. Nat Methods 8, 745–752.

